# Targeting stromal remodeling and cancer stem cell plasticity to overcome chemoresistance in triple negative breast cancer

**DOI:** 10.1101/215954

**Authors:** Aurélie S. Cazet, Mun N. Hui, Benjamin L. Elsworth, Sunny Z. Wu, Daniel Roden, Chia-Ling Chan, Joanna N. Skhinas, Raphaël Collot, Jessica Yang, Kate Harvey, M. Zahied Johan, Caroline Cooper, Radhika Nair, David Herrmann, Andrea McFarland, Nian-Tao Deng, Manuel Ruiz-Borrego, Federico Rojo, José M. Trigo, Susana Bezares, Rosalía Caballero, Elgene Lim, Paul Timpson, Sandra O’Toole, D. Neil Watkins, Thomas R. Cox, Michael Samuel, Miguel Martín, Alexander Swarbrick

## Abstract

The cellular and molecular basis of stromal cell recruitment, activation and crosstalk in carcinomas is poorly understood, limiting the development of targeted anti-stromal therapies. In mouse models of triple negative breast cancer (TNBC), Hh ligand produced by neoplastic cells reprogrammed cancer-associated fibroblast (CAF) gene expression, driving tumor growth and metastasis. Hh-activated CAFs upregulated expression of FGF5 and production of fibrillar collagen, leading to FGFR and FAK activation in adjacent neoplastic cells, which then acquired a stem-like, drug-resistant phenotype. Treatment with smoothened inhibitors (SMOi) reversed these phenotypes. Stromal treatment of TNBC patient-derived xenograft (PDX) models with SMOi downregulated the expression of cancer stem cell markers and sensitized tumors to docetaxel, leading to markedly improved survival and reduced metastatic burden. In the phase I clinical trial EDALINE, 3 of 12 patients with metastatic TNBC derived clinical benefit from combination therapy with the SMOi Sonidegib and docetaxel chemotherapy, with one patient experiencing a complete response. Markers of pathway activity correlated with response. These studies identify Hh signaling to CAFs as a novel mediator of cancer stem cell plasticity and an exciting new therapeutic target in TNBC.

**SIGNIFICANCE:** Compared to other breast cancer subtypes, TNBCs are associated with significantly worse patient outcomes. Standard of care systemic treatment for patients with non-BRCA1/2 positive TNBC is cytotoxic chemotherapy. However, the failure of 70% of treated TNBCs to attain complete pathological response reflects the relative chemoresistance of these tumors. New therapeutic strategies are needed to improve patient survival and quality of life. Here, we provide new insights into the dynamic interactions between heterotypic cells within a tumor. Specifically, we establish the mechanisms by which CAFs define cancer cell phenotype and demonstrate that the bidirectional CAF-cancer cell crosstalk can be successfully targeted in mice and humans using anti-stromal therapy.

## INTRODUCTION

Carcinogenesis draws many parallels with developmental biology. During development, dynamic interaction between stromal and epithelial cells drives patterning and function. Cell fate specification occurs through activation of transcriptional cascades in response to extracellular signals from developmental signaling pathways such as Hedgehog (Hh), Wnt, Notch, BMP (bone morphogenetic proteins) and FGF (fibroblast growth factor) (1,2). These pathways direct developmental processes either by direct cell-to-cell contact or through secreted diffusible factors (paracrine signaling). They can act individually or in concert with each other. For example, the interaction between Hh and FGF signaling pathways has been shown to mediate tracheal and lung branching morphogenesis (3,4). In mature, differentiated tissues, these pathways are quiescent but may be reactivated to drive repair and regeneration to maintain tissue homeostasis.

More specifically, the Hh developmental pathway is reactivated in a subset of cancers. Binding of Hh ligand to its receptor Patched (PTCH) enables Smoothened (SMO)-mediated translocation of Gli1 into the cell nucleus to drive the transcription of Hh target genes (5). Mutations in Hh pathway components are oncogenic drivers in “Gorlin’s-like” cancers such as medulloblastoma and basal cell carcinoma (BCC), where tumors rely on cell-autonomous Hh signaling (6). Small molecule inhibitors of SMO, Vismodegib and Sonidegib, are well tolerated and clinically approved for the treatment of these lesions (6,7). In contrast, many other solid tumors, including breast cancer, predominantly exhibit ligand-dependent pathway activation (5,6,8). While Hh signaling is quiescent in the adult mammary gland, Hh ligand expression is reactivated in a subset of breast cancers, particularly the poor-prognosis TNBC subtype (8). Breast cancer patients with a paracrine Hh pathway signature, defined by high epithelial SHH ligand expression in combination with high stromal GLI1 expression, have a greater risk of metastasis and breast cancer specific death (8).

Neoplastic cells co-opt components of the tumor microenvironment (TME) to further their progression. The TME is a complex ecosystem comprising a myriad of neoplastic and non-malignant cells embedded in a glycoprotein-rich extracellular matrix (ECM). Prominent cell types include the endothelium, cells of the immune system and cancer-associated fibroblast (CAFs). In addition to its role as a physical scaffold to support tissue architecture, the ECM also functions as a signal transducer between the different TME cell types (9). The stiffness of the ECM and the abundance of fibrillar collagen immediately adjacent to epithelial lesions provide mechanical signals that facilitate tumor development and progression (10-12). Not surprisingly, the TME has emerged as a major determinant of cancer phenotype. In breast cancer, stromal metagenes, in particular those associated with ECM remodeling, strongly predict prognosis and response to chemotherapy (13,14).

Whilst it is now apparent that Hh signals in a paracrine manner in animal models of TNBC (8) and in isolated cancer stem cells (CSCs) (15), a detailed study of the dynamic crosstalk within the TME is required to make clinical progress in integrating anti-stromal therapies into breast cancer treatment. Progress has been impeded by the field’s limited understanding of the mechanisms underlying tumor-stromal interactions, a limited repertoire of well-tolerated agents to target the TME, and an absence of predictive biomarkers for response to TME-directed therapies (16). In this study, we investigated whether, and how, SMO inhibitors (SMOi) could be used for therapeutic reprograming of the TME in human TNBC.

## RESULTS

### Hh-regulated epithelial-stromal crosstalk mediates a reversible breast cancer stem-like phenotype

To investigate the mechanistic basis for Hh-dependent tumor growth and metastasis in TNBC, we used the murine M6 allograft model of low grade TNBC, in which transgenic Hh expression drives invasion, metastasis and high-grade morphology (8) (M6-Hh; **Supplementary Fig. S1A**). Treatment with the SMO inhibitor (SMOi) GDC-0449 (Vismodegib) slowed tumor growth, reduced metastatic burden and improved overall animal survival of M6-Hh tumors (**Supplementary Fig. S1B-D**). M6-Ctrl and M6-Hh monoculture cell viability were similar between vehicle and SMOi treatment and the expression of canonical Hh target genes *Ptch*, *Gli1* and *Hhip* was downregulated *in vivo* but not *in vitro*, consistent with a paracrine requirement for Hh signaling as previously reported (8,15,17,18) (**Supplementary Fig. S1E-G**). The effects of SMO inhibition on tumor growth and gene expression were not observed in control tumors lacking Hh expression (M6-Ctrl) or in benign adult mouse mammary gland (**Supplementary Fig. S1B,G,H**), reflecting on-target drug activities. Similar results were observed with the SMOi NVP-LDE-225 (Sonidegib; **Supplementary Fig. S1I,J**).

To examine the transcriptional changes induced by Hh pathway activation in detail, we dissociated and analyzed freshly flow cytometry-sorted stromal and epithelial fractions of M6-Ctrl and M6-Hh tumors ± SMOi using RNA-Sequencing (RNA-Seq) (**Fig. 1A**). Differential gene expression analysis confirmed effective cellular fractionation, with Hh transgene expression restricted to the epithelial cell population while Hh target genes *Ptch*, *Gli1* and *Hhip* expression were induced solely in the stromal fraction of M6-Hh tumors (**Supplementary Table S1**).

**Figure 1.**
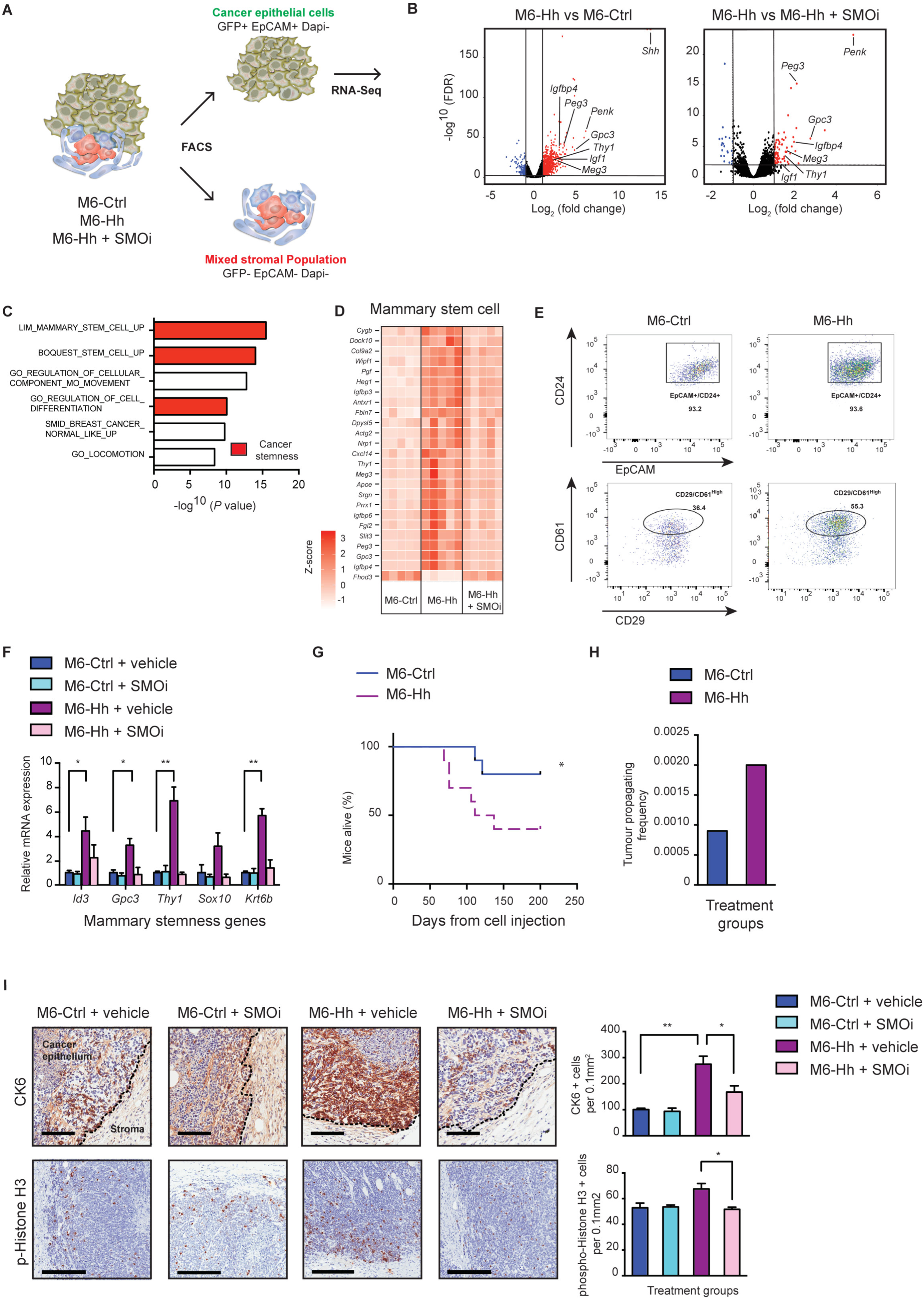
Malignant cells with increased self-renewal properties are located in the tumor epithelium adjacent to the stroma of Hh-expressing cancers. **A.** Scheme depicting the purification of epithelial and mixed stromal cell populations from disaggregated M6 murine tumor models. **B.** Expression of genes significantly downregulated (blue) or upregulated (red) (cut-off log_2_ (Fold Change) ≥ 2; vertical lines) in the cancer epithelium of M6-Hh tumors in comparison to the epithelium of M6-Ctrl or M6-Hh tumors treated with the SMO inhibitor, GDC-0449 (SMOi), plotted against FDR values (horizontal lines indicate -log^10^ (FDR) > 2). Expression level was defined by RNA-Seq analysis; each symbol represents the transcriptome of the M6 primary cancer cells from 5 biological replicates per treatment group. **C.** RNA-Seq analysis reveals significant enrichment of GSEA groups related to stemness and invasion in M6-Hh primary cells (stemness-related processes highlighted in red; FDR q-value < 0.05). **D.** Heat map showing relative expression levels of cancer stem cell genes (LIM_Mammary_Stem_Cell) in the tumor epithelium of M6-Hh primary tumors compared to the epithelium of M6-Ctrl and M6-Hh tumors treated with SMOi. Data shows normalized row Z-score (*n* = 5 biological replicates for each treatment group). **E.** Representative FACS dot plots showing the expression of cancer stem cell markers CD61 and CD29 within the EpCAM^+^/GFP^+^/CD24^+^ population of M6-Ctrl and M6-Hh tumors (*n* = 3 biological replicates per group). **F.** Relative expression of key stemness genes in M6-Ctrl and M6-Hh tumors ± SMOi. Bars represent mean ± s.e.m; *n* = 3 biological replicates per treatment group; statistical significance was determined using unpaired two-tailed Student’s t-test with equal s.d; * *P* < 0.05; ** *P* < 0.01. **G.** Kaplan-Meier curves of overall survival in mice injected with 250 primary M6-Ctrl (blue) or M6-Hh cells (violet); *n* = 10 biological replicates. Statistical significance was determined using log-rank test; * *P* < 0.05. **H.** Primary M6-Ctrl and M6-Hh tumor cells were isolated by FACS and transplanted at various dilutions into recipient mice. Limiting dilution analysis demonstrating higher *in vivo* tumor-forming capacity in M6-Hh primary cells compared to M6-Ctrl cells. *n* = 10 mice per condition. **I.** CSC plasticity at the tumor-stromal interface. Representative images showing CK6 and phospho-Histone H3 positive cancer cells at the tumor-stromal interface of M6 tumors. Scale bars: 100 µm for CK6 and 200 µm for phospho-Histone H3. Quantification of CK6-progenitor and phospho-Histone H3 positive cells at the tumor-stromal interface. Bars represent mean ± s.e.m; *n* = 3 biological replicates per treatment group. Statistical significance was determined using unpaired two-tailed Student’s t-test; * *P* < 0.05; ** *P* < 0.01.

In the epithelial compartment, 67 genes were differentially expressed (> 2-fold change, *P* < 0.001), with 60 upregulated and 7 downregulated genes in the neoplastic M6-Hh cells in comparison to M6-Ctrl or M6-Hh cells treated with SMOi. Transcriptional changes were robust and highly statistically significant (**Fig. 1B**). Hh expression in M6 cancer cells resulted in increased expression of stemness genes including *Peg3* (> 11-fold), *Igfbp4* (> 7-fold) and *Thy1* (> 4-fold) (19), which were downregulated following SMO inhibition (**Fig. 1B**).

Gene Set Enrichment Analysis (GSEA) and Gene Ontology (GO) analysis of the purified epithelial fraction highlighted enrichment for genes specifically and almost exclusively associated with mammary stemness and invasion, consistent with the morphologically undifferentiated phenotype previously observed in Hhoverexpressing tumors **(8)** (**Fig. 1C,D** and **Supplementary Table S1**). To examine the cancer stem cell-like (CSC) phenotype in greater detail, epithelial cells were profiled by flow cytometry. M6-Hh tumors had a higher proportion of CD61^hi^ cells, previously shown to be a marker of mouse mammary CSCs (20), within the EpCAM^+^CD24^hi^CD29^+^ cancer cell population (55.3% in M6-Hh tumors vs 36.4% in M6-Crtl tumors; **Fig. 1E**). M6-Hh tumors also had elevated expression of the stemness markers *Id3, Gpc3, Thy1, Sox10 and Krt6* (19-22), validating the RNA-Seq data (**Fig. 1B,F**). Following transplantation of low numbers of sorted primary M6-Hh and M6-Ctrl cells into naive recipients, tumor latency was shorter and penetrance markedly higher in the M6-Hh group (**Fig. 1G**). Limiting dilution assays (23) were used to quantitate the impact of Hh signaling on tumor-initiating capacity. M6-Ctrl and M6-Hh tumor cells were isolated by FACS and transplanted at various dilutions. M6-Hh cells had significantly higher tumor-initiating capacity (1 in 435) compared with M6-Ctrl cells (1 in 1088; **Fig. 1H**). Importantly, the proliferation and expression of CSC markers were indistinguishable between M6-Ctrl and M6-Hh cells in monoculture, indicating that Hh expression in M6 cells does not regulate CSC properties in a cell autonomous manner (**Supplementary Fig. S1E**).

Immunohistochemical detection of the mammary progenitor marker cytokeratin 6 (CK6; product of the *Krt6* gene) (24) localized cells with a stem/progenitor signature specifically to the tumor-stromal interface (**Fig. 1I**). SMO inhibition reduced the expression of *Id3, Gpc3, Thy1, Sox10 and Krt6* and significantly reduced the number of cells positive for CK6 and the mitotic marker phospho-Histone H3 at the tumor-stromal interface (**Fig. 1F,I**). These data demonstrate that paracrine Hh signaling results in the induction of a reversible stem-like phenotype preferentially at the tumor-stromal interface.

### Stromal Hh signaling leads to marked ECM-related gene expression changes and associates with poor prognosis in patients with TNBC

RNA-Seq analysis of the stromal fraction revealed 185 genes that were differentially expressed (> 2-fold, *P* < 0.001), with 146 upregulated and 39 downregulated genes in the stroma of M6-Hh tumors compared to M6-Ctrl and M6-Hh tumor + SMOi (**Supplementary Table S1**). A number of genes were markedly upregulated by Hh signaling, in particular the growth factor gene *Fgf5* at more than 290-fold, *St8Sia2* (> 40-fold) and *Tspan11* (> 4-fold; **Fig. 2A**). The large majority of gene expression changes in Hh-activated stroma returned to baseline following treatment with SMOi (**Fig. 2A** and **Supplementary Table S1**), suggesting that the stromal transcriptional changes are SMO-dependent and reversible.

**Figure 2.**
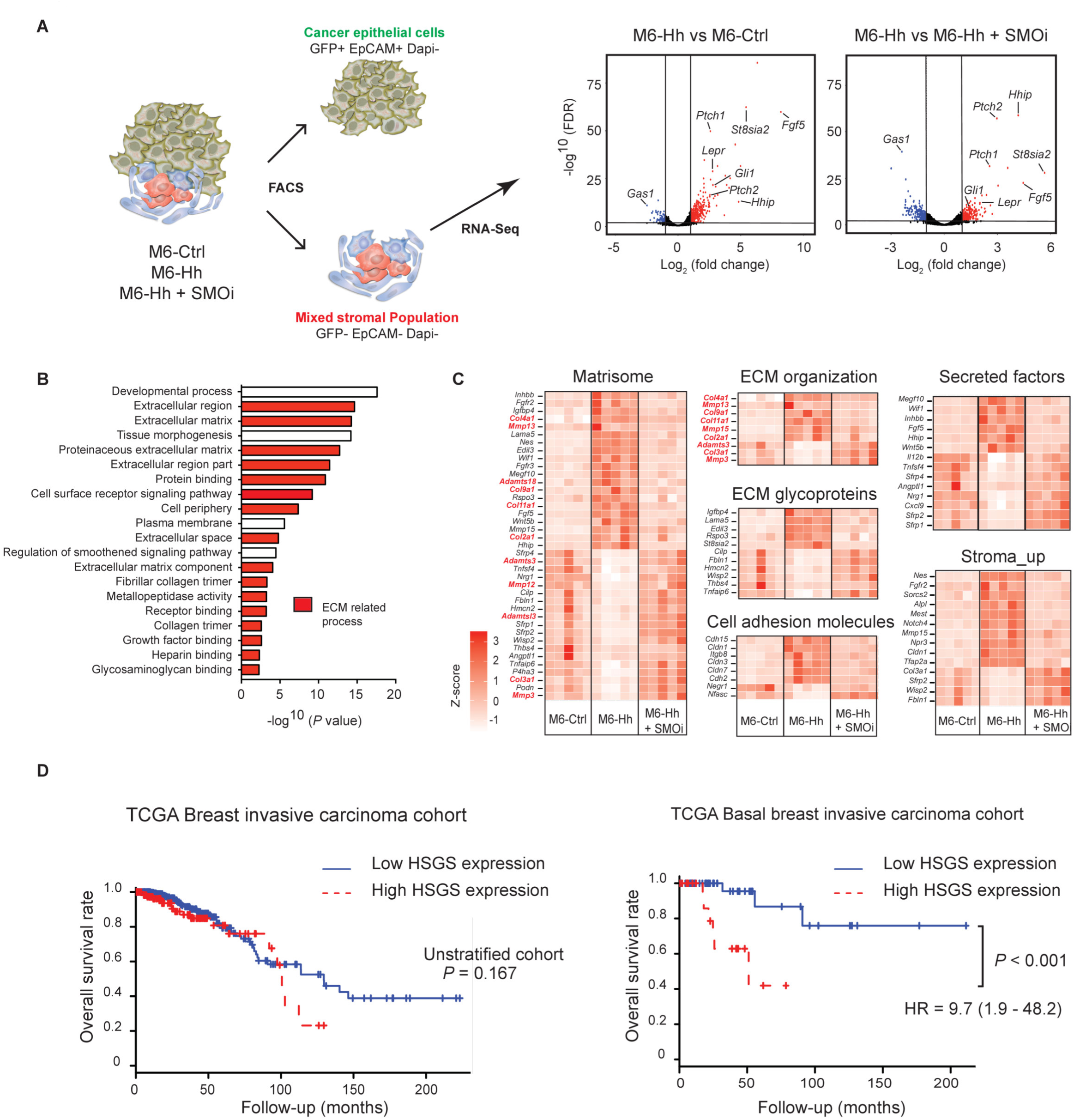
Hh-activated stroma defines a novel transcriptional signature robustly associated with poor-prognosis in TNBC. **A.** Expression of genes significantly downregulated (blue) or upregulated (red) (cutoff log_2_ (Fold Change) ≥ 2; vertical lines) in the stroma of M6-Hh tumors compared to the stroma of M6-Ctrl or M6-Hh tumors treated with SMOi, plotted against FDR values (horizontal lines indicate -log^10^ (FDR) > 2). Each symbol represents the transcriptome of the stromal population from 5 biological replicates per treatment group. **B.** RNA-Seq analysis reveals significant enrichment of GO groups related to ECM organization and production in M6-Hh tumor stroma (ECM-related processes highlighted in red; FDR q-value < 0.05). **C.** Heat maps demonstrating relative expression levels of gene sets defined by RNA-Seq analysis. Genes highlighted in red represent those that are directly involved in ECM production. Data shows normalized row Z-score (*n* = 5 biological replicates for each treatment group). **D.** Kaplan-Meier curves of overall survival in unstratified breast cancer patients or in patients with basal, TNBC. Blue and red lines represent low and high Hh-stromal gene signature expression (HSGS), respectively. Statistical significance was determined using the Log-rank test; *** *P* < 0.001.

Comparative GO and GSEA analysis revealed ECM processes as highly enriched in the stromal fraction of M6-Hh tumors (**Fig. 2B,C**), suggesting that a major influence of paracrine Hh activation on the tumor stroma is related to ECM production and remodeling (25). Genes highly enriched in this set included collagens (*Col2a1 Col3a1, Col4a1*, *Col9a1*, *Col11a1*), ECM-remodeling metallopeptidases (*Mmp3, Mmp13*, *Mmp15*, *Adamts3, Adamts18*), ECM glycoproteins (*St8sia2*, *Rspo3*, *Lama5*, *Edil3*, *Thbs4*) and cell adhesion molecules (*Cldn3*, *Cldn7*, *Cldn1*, *Cdh2*, *Cdh15*) (**Fig. 2C**). Interestingly, these ECM gene expression changes were not a result of changes in stromal cellular composition due to Hh pathway activation and long-term SMO inhibition. Immunohistochemical analysis did not reveal any difference between M6-Ctrl and M6-Hh tumors in terms of CAFs, endothelial and innate immune cell abundance, regardless of treatment with SMOi (**Supplementary Fig. S2A**). This finding was confirmed using whole tumor qRT-PCR and flow cytometry analysis (**Supplementary Fig. S2B,C**).

To determine the prognostic value of the Hh-activated stromal gene signature (HSGS), we examined its impact on overall survival using The Cancer Genome Atlas (TCGA) breast invasive carcinoma cohort. The HSGS was not predictive of patient outcome in the unstratified patient cohort (**Fig. 2D**) but was associated with significantly lower patient overall survival uniquely in the basal breast cancer subtype, where Hh ligand is most frequently overexpressed (8) (Hazard ratio = 9.7 (1.9 - 48.2); *P* < 0.001; **Fig. 2D**).

Accumulating evidence suggests that CAFs contribute to tumor growth upon Hh ligand activation (15,26). However, the stroma of M6 tumors is composed of multiple cell types, any of which may be responsible for Hh-dependent gene expression changes. We used a single-cell RNA sequencing (scRNA-Seq) approach to determine the cell population/s responding to paracrine Hh signaling. The microfluidic 10X Chromium system was used to comprehensively profile gene expression at cellular resolution in thousands of cells isolated from freshly dissociated M6-Ctrl and M6-Hh tumors ± SMOi. In total, we compared 6,064 FACS-isolated cells from M6-Ctrl tumors, 6,200 cells from M6-Hh tumors, and 2,686 single cells from M6-Hh tumor treated with SMOi.

As shown in **Figure 3A**, unsupervised clustering analysis of 14,950 cells revealed populations of myeloid, neoplastic, endothelial, CAF and natural killer cells within the breast TME (**Fig. 3A**). Importantly, the upregulation of canonical Hh target genes *Gli1*, *Ptch1*, *Ptch2* and *Hhip* and ECM genes such as *Col4a1*, *Tspan11*, *St8sia2* and *Fgf5* was observed exclusively in the CAF population of M6-Hh compared to M6-Ctrl tumors (**Fig. 3A**), and not in other stromal cell types. More specifically, the ECM signature detected in the stroma of Hh-expressing tumors *via* ‘bulk’ RNA-Seq was driven by CAF gene expression (**Fig. 2B,C**; **Fig. 3B**). This scRNA-Seq approach also confirmed the lack of autocrine Hh pathway activation within the neoplastic cells (**Fig. 3A**). Treatment with SMOi almost completely reversed the Hh-dependent gene expression changes observed in CAFs without affecting gene expression in other stromal cell types (**Fig. 3A**; **Supplementary Fig. 2D** and **Supplementary Table S2**), highlighting the on-target activity of SMOi at the single cell level.

**Figure 3.**
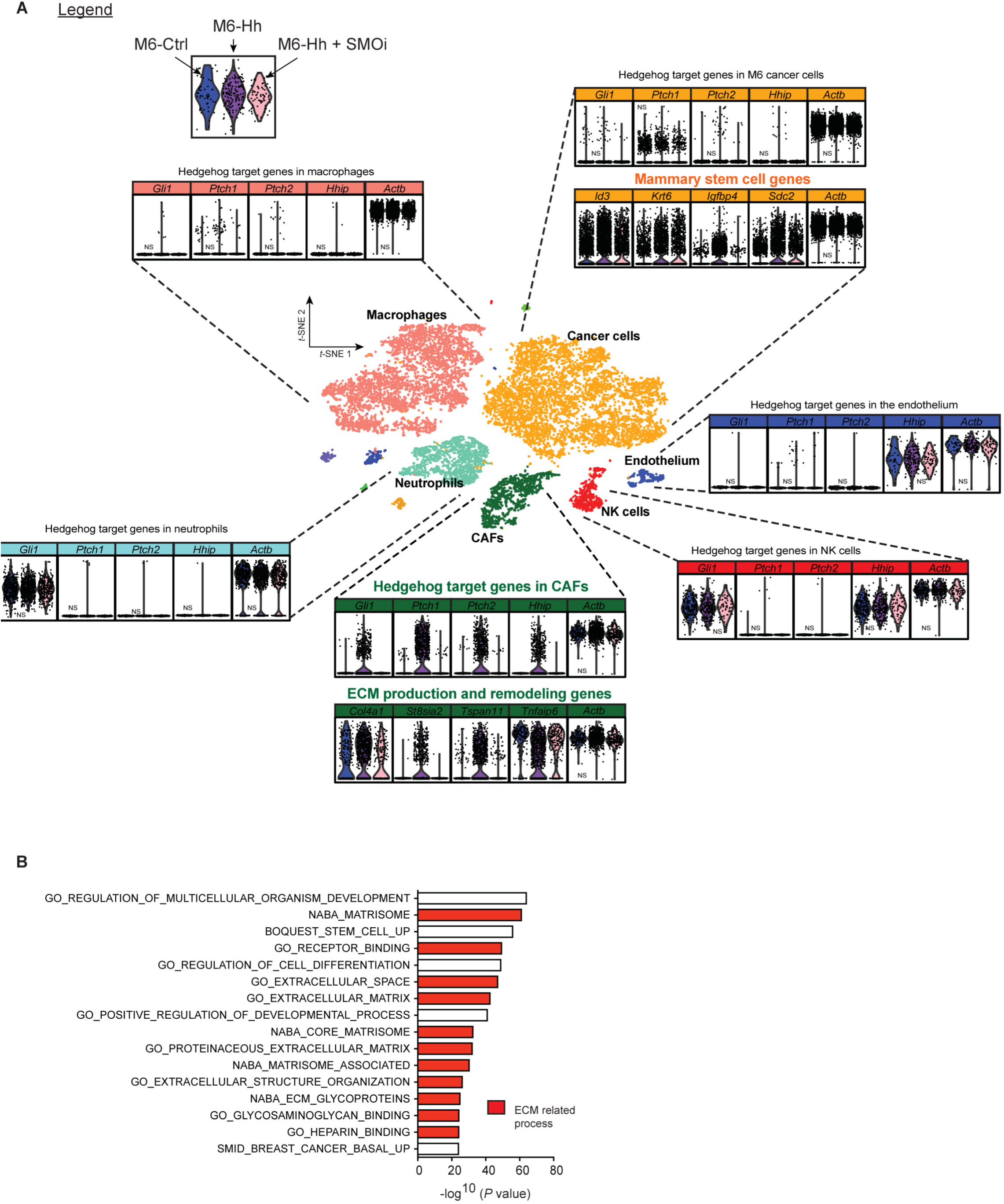
Paracrine Hedgehog signaling at cellular resolution. **A.** Freshly isolated M6-Ctrl (blue), M6-Hh (magenta) and M6-Hh tumors + SMOi (pink violin plot) were captured using the 10X Chromium technology and the Cell Ranger Single Cell Software Suite 2.0 was used to perform demultiplexing, barcode processing, and single cell 3′gene counting. *t*-SNE plot shows the subcellular clusters present in the breast tumor microenvironment of M6 tumors. Unless stated, *P*-values are highly significant (**Supplementary Table S2**). **B.** Single cell RNA-Seq analysis reveals significant enrichment of GSEA groups related to ECM organization and production specifically in the CAF population of M6-Hh tumors (ECM-related processes highlighted in red; FDR q-value < 0.05).

Co-culture of primary CAFs with M6-Hh cells was sufficient to recapitulate this dynamic epithelial-stromal crosstalk resulting in the induction of Hh target gene expression in the CAFs (**Supplementary Fig. S3A,B**) and concomitant upregulation of CSC markers *Id3, Gpc3, Itgb3* and *Krt6b* in M6-Hh cells compared to M6-Ctrl + CAF co-culture systems (**Supplementary Fig. S3A,C**). Importantly, this stromal-epithelial malignant crosstalk was blocked by SMOi (**Fig. 3A**, **Supplementary Fig. S3B,C**). These data allow us to conclude that Hh signaling occurs solely in a paracrine manner in this murine model of TNBC and CAFs are the therapeutic target of SMOi in TNBC.

### Hh activated CAFs drive fibrillar collagen deposition and remodeling, resulting in mechanosignaling and a stem-like phenotype in adjacent neoplastic cells

Bulk and scRNA-Seq data suggested that stromal Hh-signaling drives collagen remodeling in the local ECM (**Fig. 2B,C** **and** **Fig. 3**), which is known to associate with breast cancer progression (27,28). We employed second harmonic generation (SHG) microscopy, a sensitive label-free method for quantifying fibrillar collagen density and orientation in tissues (29). SHG analysis revealed a ~3-fold increase in fibrillar collagen density at the tumor stromal-interface of Hh-expressing tumors (**Fig. 4A**), but not in the tumor center (data not shown). The increase in collagen abundance was confirmed by chromogenic staining using Picrosirius red (**Fig. 4B**). Further detailed analyses of the distribution and orientation of collagen fibers as described by Mayorca-Guiliani *et al*. (30) and by Gray level co-occurrence matrix (GLCM) analysis (31,32) revealed changes in texture and cross-linking of the ECM with linearization of collagen fibers adjacent to epithelial lesions in M6-Hh tumors, a hallmark of breast tumor growth and invasiveness (10,30) (**Fig. 4A**). These features of the collagen ECM in Hh-expressing tumors were ameliorated in mice treated with SMOi (**Fig. 4A**), demonstrating an ongoing dependency on SMO activation.

**Figure 4.**
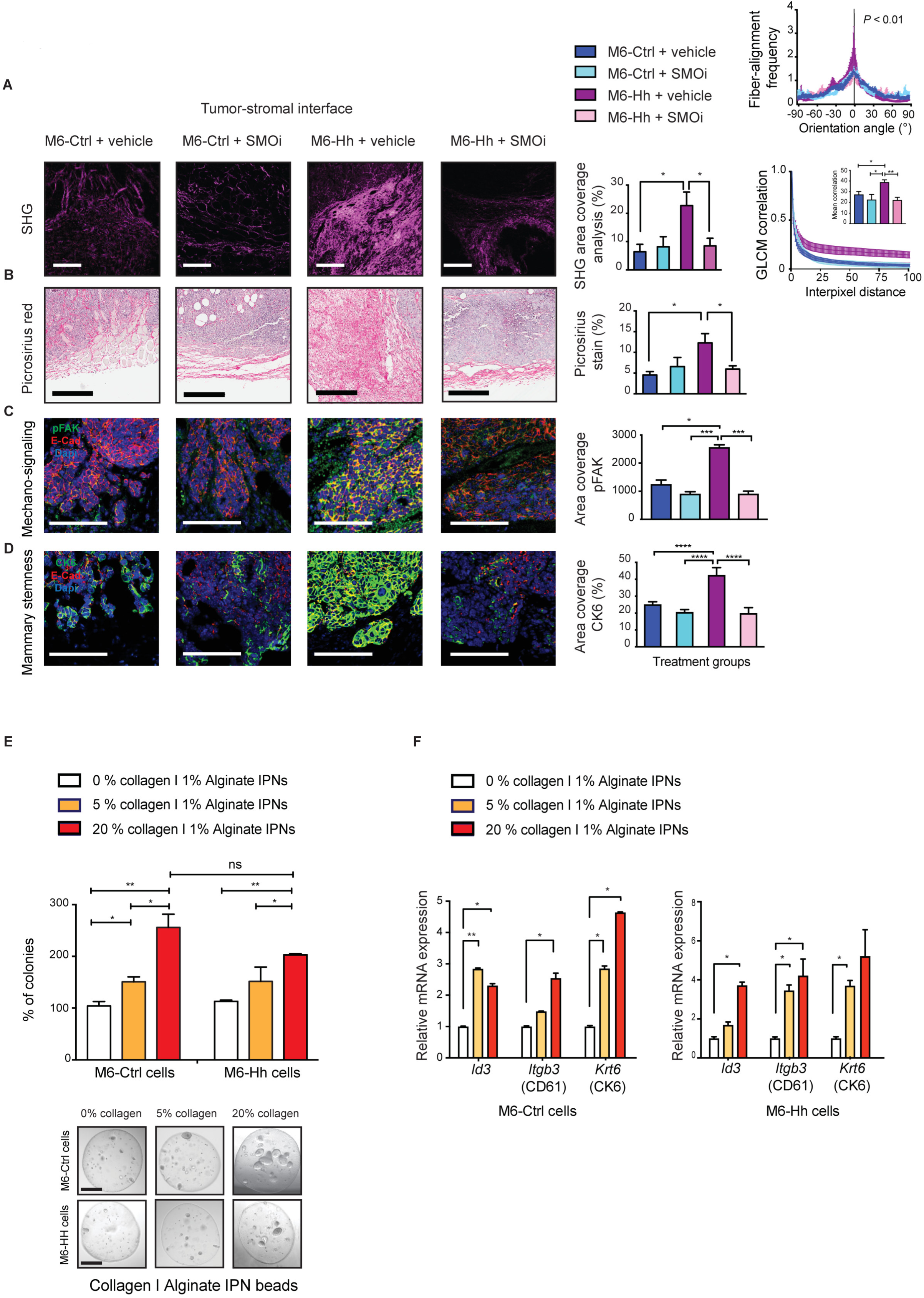
High fibrillar collagen content dependent on paracrine Hh pathway activation promotes high mechanosignaling and breast cancer stemness. **A-D.** Concomitant expression analysis of collagen content and organization, integrin/focal adhesion activation and CSC-like characteristics of M6 tumor models ± SMOi (*n* = 3 biological replicates). **A.** Representative multi-photon SHG imaging (scale bars: 100 µm) and quantitative analysis of collagen abundance at the tumor-stromal interface. Bars represent mean ± s.e.m; statistical significance was determined using unpaired two-tailed Student’s t-test with equal s.d. * *P* < 0.05. Corresponding graphs comparing fiber orientation (top right panel) and quantifying gray level cooccurrence matrix (bottom right panel) in the different M6 models. Bars represent mean correlation distance ± s.e.m. The unpaired two-tailed nonparametric Mann– Whitney U test was used for determining statistical significance across distributions. ** *P* < 0.01. For the GLCM analysis, statistical significance was determined using unpaired two-tailed Student’s t-test with equal s.d; * *P* < 0.05; ** *P* < 0.01. **B.** Collagen I and III deposition at the tumor-stromal interface detected and quantified by picrosirius red staining. Scale bars, 200 µm. Bars represent mean ± s.e.m. Statistical significance was determined using unpaired two-tailed Student’s t-test with equal s.d; * *P* < 0.05. **C.** Representative immunofluorescence images and quantification of phospho-FAK and **D.** the CK6 progenitor expression at the tumor-stromal interface of M6 tumor models. Scale bars, 100 µm. Statistical significance was determined using Kruskal-Wallis test; * *P* < 0.05; *** *P* < 0.001; **** *P* < 0.0001. **E.** M6-Ctrl and M6-Hh single cells were embedded within and grown for 12 days in 3D spherical 1% Alginate IPNs containing increasing concentrations of collagen (0% to 20%). Quantification of the number of colonies formed was normalized to the mean colony count in 0 % collagen IPNs. Bars represent mean ± s.e.m; *n* = 3 biological replicates with at least 6 technical replicates per condition. Statistical significance was determined using unpaired two-tailed Student’s t-test with equal s.d; * *P* < 0.05; ** *P* < 0.01. Representative phase contrast images of M6 colonies on polyacrylamide substrata after 12 days of culture. Scale bars: 100 µm. **F.** Relative mRNA expression of the CSC markers *Id3*, *Igtb3* and *Krt6b* in M6-Ctrl and M6-Hh cells cultured within 3D alginate-collagen I IPNs. Bars represent mean ± s.e.m; *n* = 3 biological replicates with 6 technical replicates per experiment. Statistical significance was determined using unpaired two-tailed Student’s t-test with equal s.d. * *P* < 0.05; ** *P* < 0.01.

Sites of collagen deposition and cross-linking at the stromal-epithelial interface were also marked by increased phosphorylation and activation of focal adhesion kinase (FAK) on cancer cells, a key signaling intermediate downstream of integrin receptors (**Fig. 4C**). These cells also expressed elevated cytokeratin 6 (**Fig. 4D**), correlating fibrillar collagen content to FAK signaling and the acquisition of a stem-like phenotype in the neoplastic cells. Importantly, mechanosignaling and cancer stemness exclusively occurred in close proximity to rich dense collagen regions and were not observed in the core of the M6-Hh tumors (**Supplementary Fig. S4A,B**).

To directly assess the sufficiency of collagen mechanosignaling to promote stemness, the clonogenic potential of M6-Ctrl and M6-Hh cells was assessed using 3-dimensional cultures encapsulated within Alginate-Collagen I Inter-Penetrating Network (IPN) hydrogels. The enrichment for fibrillar collagen in this *in vitro* model recapitulates the features of stromal collagen matrix deposition observed in Hh-expressing models. Increased content and presence of highly bundled fibrillar collagen significantly increased the clonogenic capacity of M6 cells, a functional surrogate for CSC activity (33), independently of Hh ligand expression (**Fig. 4E**). Furthermore, increasing collagen I abundance also increased expression of the stem cell markers *Id3*, *Itgb3* (CD61) and *Krt6* (CK6) (**Fig. 4F**). These data demonstrate that Hh-dependent stromal ECM remodeling is sufficient to foster a CSC phenotype.

### Paracrine Hh-FGF5 signaling also contributes to CSC plasticity

To identify additional mechanisms by which stromal signaling promotes the acquisition of a CSC phenotype, we turned our attention to *Fgf5,* which was strongly upregulated in Hh-activated stroma compared to controls (**Fig. 2A,C** and **Supplementary Table S1 and S2**). qRT-PCR analysis of whole tumors confirmed ~60-fold upregulation of *Fgf5* mRNA in M6-Hh tumors, which was reversed upon SMOi treatment (**Fig. 5A**). Notably, a subset of Hh-activated CAFs exhibited robust expression of *Fgf5* at the single-cell resolution, reflecting the spatial localization of these CAFs, in close proximity with M6-Hh cells (**Fig. 5B**). Immunohistochemical analysis of phospho-FGFR revealed potent receptor activation in M6-Hh tumors, primarily in epithelial cells adjacent to stroma, which was reversed upon SMOi treatment (**Fig. 5C**). To explore the role of FGF5 in the acquisition of the CSC phenotype, M6-Ctrl cells were treated *in vitro* with recombinant FGF5 protein and proliferation, stem cell marker expression and sphere forming capacity were evaluated. FGF5 treatment led to a modest increase in proliferation under serum and growth factor deprivation (**Supplementary Fig. S5A**). In contrast, the stemness markers *Id3* and *Sox10* were robustly upregulated (**Fig. 5D**) and primary and secondary sphere forming capacity increased by ~3-fold (**Fig. 5E** and **Supplementary Fig. S5B**), suggesting a specific effect of FGF5 signaling on stemness.

**Figure 5.**
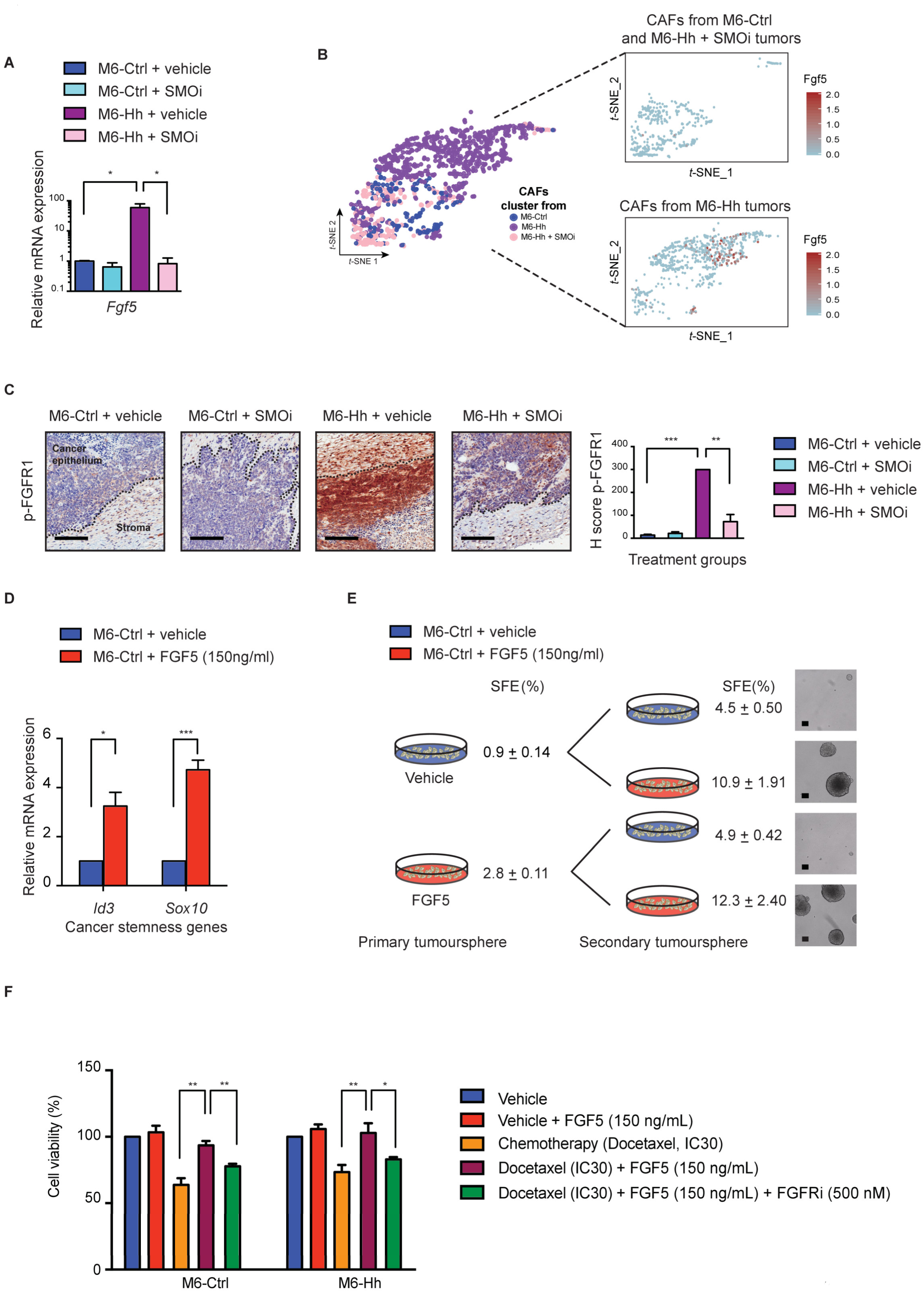
Hh-activated cancer-associated fibroblasts form a reversible, chemoresistant CSC niche *via* FGF pathway activity. **A.** RT-qPCR analysis of *Fgf5* expression in M6 whole tumors. Bars represent mean ± s.e.m; *n* = 3 biological replicates per treatment group with 3 technical replicates per assay. Statistical significance was determined using unpaired two-tailed Student’s t-test with equal s.d; * *P* < 0.05. **B.** Expression of *Fgf5* at the single cell resolution in M6 tumor models. **C.** Representative immunohistochemistry staining for phospho-FGFR on M6 tumors. Scale bars: 100 µm. Quantification of phospho-FGFR positive cancer cells at the tumor-stromal interface. Bars represent mean ± s.e.m; *n* = 4 biological replicates per treatment group; statistical significance was determined using unpaired two-tailed Student’s t-test with equal s.d; ** *P* < 0.01; *** *P* < 0.001. **D.** Relative mRNA expression of CSC markers *Id3* and *Sox10* in M6-Ctrl cells treated with DMSO (vehicle) or recombinant FGF5 *in vitro.* Bars represent mean ± s.e.m; *n* = 3 biological replicates with 3 technical replicates per experiment; statistical significance was determined using unpaired two-tailed Student’s t-test; \**P* < 0.05; *\*\*\* P* < 0.001. **E.** Primary and secondary tumorsphere formation of M6-Ctrl cells treated with DMSO (vehicle) or recombinant FGF5. Sphere Formation Efficiency (SFE) values in % are mean ± s.e.m; *n* = 3 biological replicates with 3 technical replicates per tumorsphere assay. Representative phase contrast micrographs of M6-Ctrl spheres upon recombinant FGF5 stimulation. Scale bars: 100 µm. **F.** Cell viability of M6-Ctrl and M6-Hh cells treated with indicated agents (*n* = 5 biological replicates with 6 technical replicates each). Bars represent mean ± s.e.m; statistical significance was determined using unpaired two-tailed Student’st-test with equal s.d; * *P* < 0.05; ** *P* < 0.01.

To test whether the induction of sphere forming capacity by FGF5 treatment was epigenetically stable or plastic, we tested the impact of addition or removal of FGF5 to primary and secondary sphere cultures. Increased sphere formation in response to FGF5 was observed in secondary cultures regardless of whether those cells were pre-treated with FGF5 during primary cultures (**Fig. 5E** and **Supplementary Fig. S5B)**. Furthermore, removal of recombinant FGF5 decreased secondary sphere formation to levels comparable to those of cells never treated with FGF5 ligand (**Fig. 5E** and **Supplementary Fig. S5B)**. Consistent with this result, treatment of spheres with a small molecule inhibitor of FGFR signaling (NVPBGJ398) prevented FGF5 mediated tumorsphere formation in primary or secondary cultures (**Supplementary Fig. S5C**). These results demonstrate that stromal-derived FGF5 is sufficient to promote reversible transition to a CSC phenotype, rather than through the expansion of a sub-population of CSC.

The CSC phenotype is associated with resistance to cytotoxic chemotherapy in TNBC (34). To test whether the FGF-dependent increase in CSC alters the sensitivity of TNBC cells to chemotherapy, the efficacy of docetaxel was evaluated in M6 cell lines *in vitro*. Monocultures of M6-Hh cells did not display differential sensitivity to docetaxel when compared to M6-Ctrl cells as expected (**Supplementary Fig. S5D**). However, stimulation of M6-Ctrl and M6-Hh models with recombinant FGF5 ligand rescued these cells from docetaxel cytotoxicity (**Fig. 5F**). The FGFR inhibitor NVPBGJ 398 abrogated drug resistance conferred by FGF5 (**Fig. 5F**). Similar results were observed in the human MDA-MB-231 cell line model of TNBC (**Supplementary Fig. S5E**). This result suggests that FGF5, released by Hh-activated CAFs, creates a “chemo-resistant niche” at the tumor-stromal interface that can serve as a reservoir for eventual tumor relapse in TNBC. It also suggests that targeting both tumor and stromal compartments with chemotherapeutic regimen and SMOi, respectively, may be an effectively therapeutic strategy.

### Stromal SMO inhibition combined with chemotherapy markedly improves preclinical TNBC outcomes

To directly test these findings in more clinically relevant models, we turned our analysis to xenograft models of human TNBC. All three patient-derived xenograft (PDX) models tested were Hh ligand-positive as was the MDA-MB-231 cell line model (**Supplementary Fig. S6A**). We found convincing evidence of exclusively stromal-restricted Hh signaling, using species-specific RT-PCR and sensitivity to SMOi (**Supplementary Fig. S6B,C**). In addition, *in vitro* treatment of MDA-MB-231 cells with SMOi did not alter Hh target gene expression or proliferation (**Supplementary Fig. S6D,E**), consistent with a paracrine requirement for Hh signaling in these models. The HCI-002 and MDA-MB-231 models were chosen for further investigation, as they are well-accepted models for TNBC (35,36).

Analysis of the ECM in HCI-002 xenografts by SHG and picrosirius red staining revealed abundant fibrillar collagen exclusively at the tumor-stromal interface (**Fig. 6A,B**) that was highly linearized and densely packed as depicted by orientation and GLCM analysis (**Fig. 6A**). Areas of high fibrillar collagen density were also associated with concomitant FAK phosphorylation and expression of the human CSC marker ALDH1 (**Fig. 6C**). As observed in the transgenic model, collagen density and orientation, FAK phosphorylation and CSC marker expression were reduced following SMO inhibition (**Fig 6A-C**).

**Figure 6.**
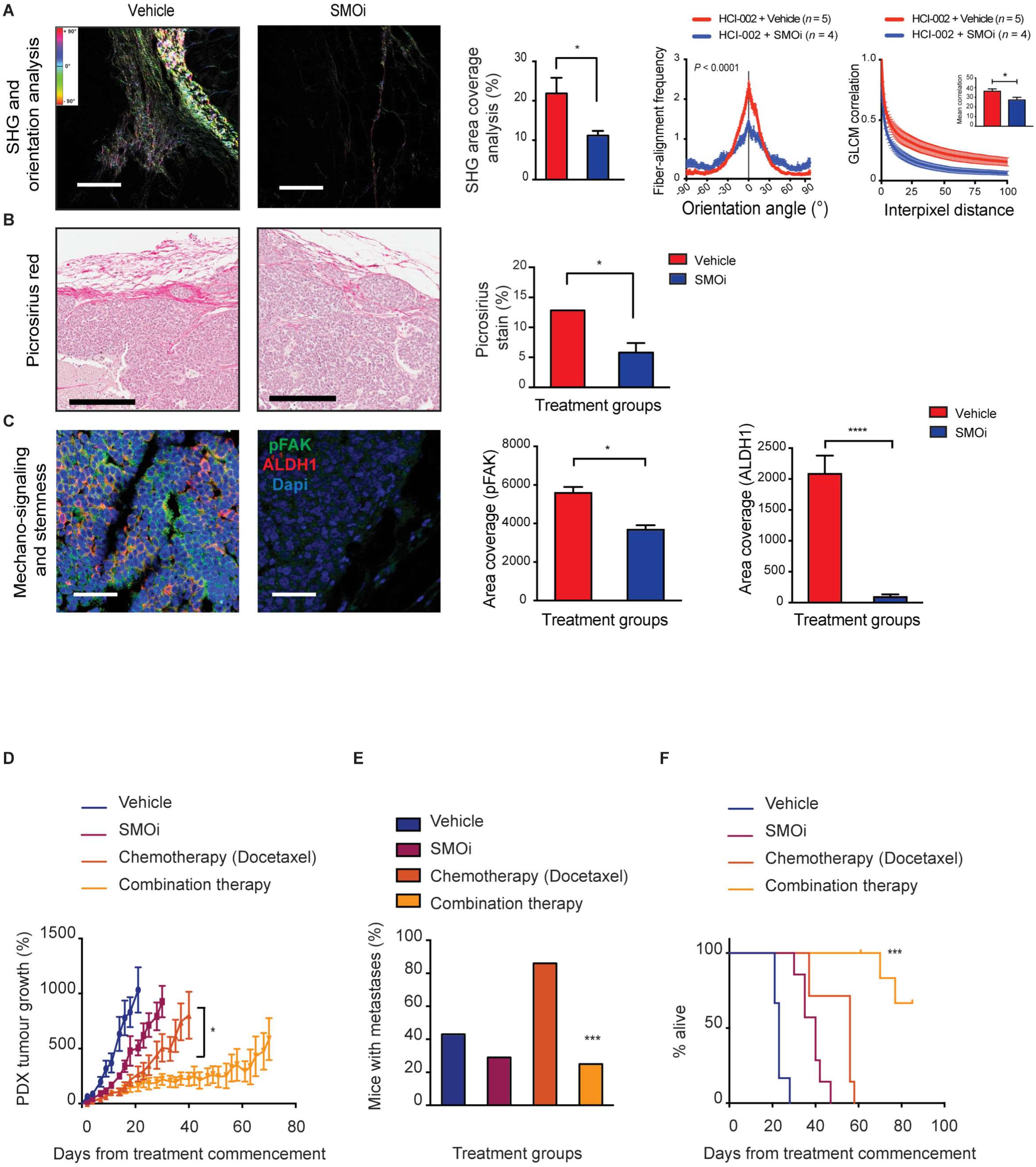
Efficacy of long-term SMO-combined therapy in preclinical models. **A-C.** Concomitant expression analysis of collagen content and organization, integrin pathway activation and CSC-like characteristics of the TNBC HCI-002 PDX model treated with vehicle (red) or the SMO inhibitor (NVP-LDE225; blue) (*n* = 5 biological replicates). **A.** Representative imaging (scale bars: 100 μm) and quantitative analysis of collagen abundance at the tumor-stromal interface (left panel). Bars represent mean ± s.e.m; statistical significance was determined using unpaired two-tailed Student’s t-test with equal s.d. * *P* < 0.05. Corresponding graphs comparing fiber orientation (middle) and quantifying gray level co-occurrence matrix analysis (right panel) in the vehicle and SMOi-treated models. The unpaired twotailed nonparametric Mann–Whitney U test was used for determining statistical significance across distributions. **B.** Collagen I and III deposition at the tumor-stromal interface detected and quantified by picrosirius red staining. Scale bars, 200 μm. Bars represent mean ± s.e.m. Statistical significance was determined using unpaired twotailed Student’s t-test with equal s.d; * *P* < 0.05. **C.** Representative immunofluorescence images and quantification of concomitant phospho-FAK (green) and the human CSC marker ALDH1 (red) expression at the tumor-stromal interface of this PDX model. Scale bars, 100 μm. Bars represent mean ± s.e.m; *n* = 3 biological replicates. Statistical significance was determined using Kruskal- Wallis test; * *P* < 0.05; **** *P* < 0.0001. **(D-F)** TNBC HCI-002 PDX model treated with vehicle (blue), SMO inhibitor (NVP-LDE225; magenta), chemotherapy (docetaxel; dark orange) or NVP-LDE225 + docetaxel (orange line) (*n* = 7 mice per treatment group). **D.** Tumor growth curves. Data represent mean ± s.e.m; statistical significance was determined using unpaired Student’s t test between combination therapy and docetaxel monotherapy; * *P* < 0.05. **E.** Percentage of mice with detectable metastases in the lung, liver and axillary lymph node in each treatment group. Statistical significance was determined using the Fisher’s exact test; *** *P* < 0.001. **F.** Kaplan-Meier curves of mice overall survival of each treatment group. Statistical significance was determined using the Log-rank test of NVP-LDE225 + docetaxel versus docetaxel; *** *P* < 0.001.

Based on these observations we predicted that SMOi would sensitize tumors to cytotoxic chemotherapy. HCI-002 PDX and MDA-MB-231 xenografts were then treated with SMOi +/- docetaxel (**Fig. 6D-F**). Compared with vehicle treatment, either SMOi or docetaxel monotherapy slowed tumor growth. However, the most robust and durable therapeutic effect occurred with combined therapy (**Fig. 6D,F**). Interestingly, the proportion of mice with metastatic disease at ethical endpoint (based on primary tumor size) was doubled in the docetaxel treated group, an observation previously made with paclitaxel in TNBC mouse models (37) (**Fig. 6E**). Combination therapy reduced the frequency of mice with metastatic disease to below that seen in the vehicle control group, despite these mice being alive much longer compared to those in the other treatment groups. Similar therapeutic benefit was observed in MDA-MB-231 tumor bearing mice treated with combination therapy in terms of tumor growth, overall survival and histological changes (**Supplementary Fig. S7A-D**).

We then examined the impact of stromal Hh pathway inhibition on the histology of the tumor epithelial and stromal compartments. The proportion and number of stromal cells in the HCI-002 PDX model were unaffected by long-term SMOi and/or chemotherapy treatments (data not shown). Even in the context of docetaxel chemotherapy, Hh signaling was still stromally-restricted (**Supplementary Fig. S7E,F**), arguing that SMOi does not sensitize cells to docetaxel *via* a cell-autonomous mechanism.

### Clinical evaluation of SMOi and docetaxel combination therapy in the phase I clinical trial EDALINE

The promising preclinical study results led us to establish the EDALINE (GEICAM/2012-12) phase I trial of docetaxel in combination with SMOi (NVP-LDE225, Sonidegib) to determine the maximum tolerated dose (MTD) and the recommended phase II dose (RP2D) of this combined therapy in patients with metastatic TNBC. Twelve patients with prior standard of care chemotherapy treatments with taxanes and/or anthracyclines were enrolled. Detailed information on clinical trial design, patient and treatment characteristics are described in **Supplementary Tables S3 and S4** and on ClinicalTrials.gov (Identifier: NCT02027376; <u>https://clinicaltrials.gov/ct2/show/NCT02027376?term=edaline&rank=1</u>). Combination therapy was well tolerated and the RP2D of Sonidegib 800 mg once daily in combination with Docetaxel 75mg/m^2^ every 21 days was established. Clinical response was evaluated according to standard clinical trial RECIST (38) (Response Evaluation Criteria in Solid Tumors) criteria version 1.1. One patient experienced a complete clinical response, defined as the disappearance of all target and non-target tumor lesions and the absence of new tumor lesions. As shown in **Figure 7A**, the 54-year old postmenopausal woman presented with metastatic disease in the lungs (red and blue arrows). Following 8 cycles of combined therapy, the patient achieved complete clinical response, evident by the complete resolution of all her lung metastases on routine progress CT scan (**Fig. 7A** and **Supplementary Tables S3 and S4**). Two other patients experienced disease stabilization (**Supplementary Table S3**).

**Figure 7.**
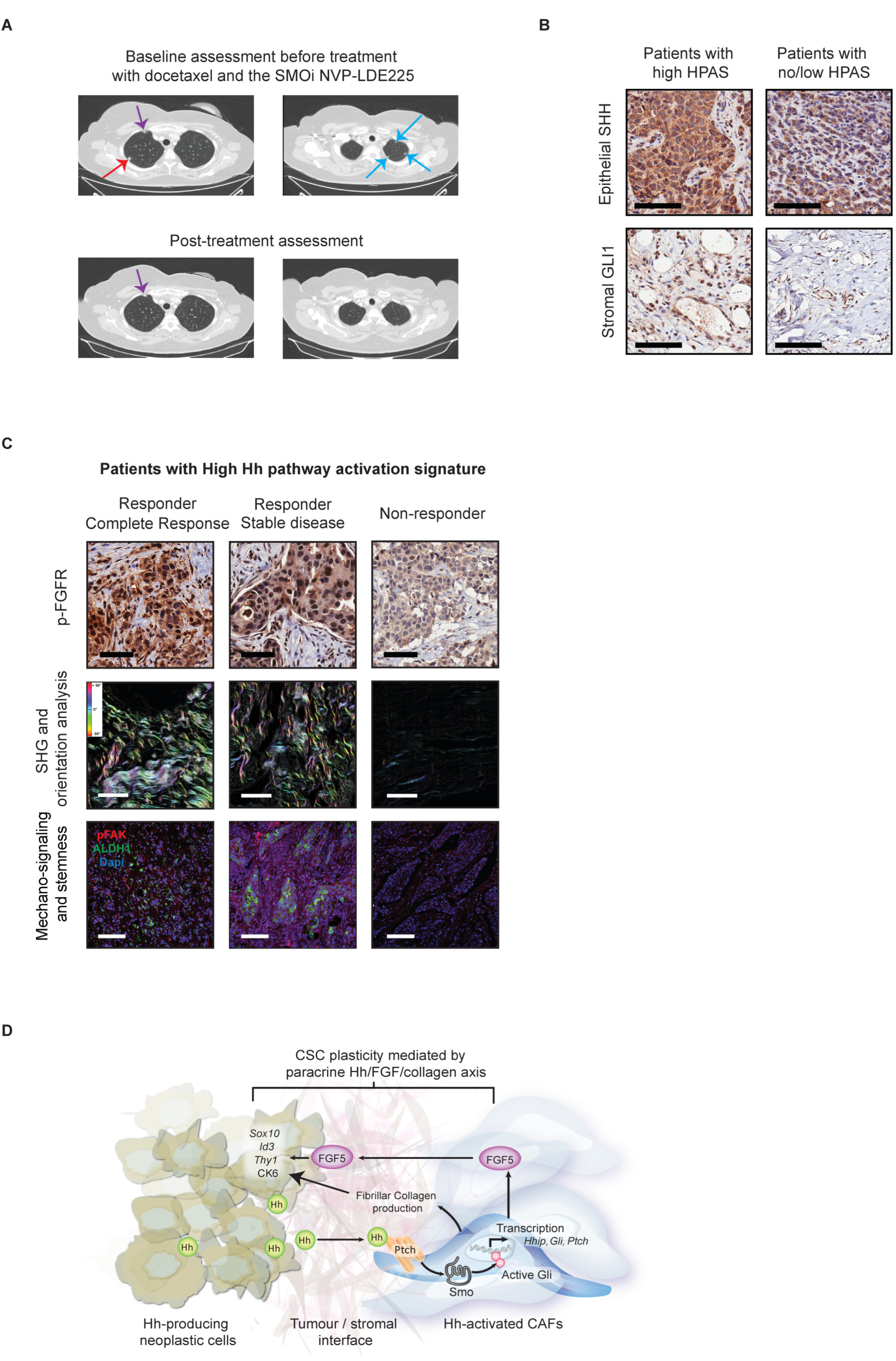
Phase I clinical trial of docetaxel and SMO inhibitor, NVP-LDE225 (EDALINE) in patients with advanced TNBC. **A.** Representative Computed Tomography (CT) images from a patient with complete radiological response. The 54-year old postmenopausal woman was diagnosed with recurrent metastatic TNBC in the lungs with one measurable lesion on the right upper lobe (red arrow) and several non-measurable lesions (blue arrows). Therapeutic response was evaluated according to RECIST criteria version 1.1. The remained image seen in the upper part of the upper right lobe corresponds to the azygos vein (violet arrow). **B.** Representative SHH and GLI1 immunostaining of naive tumor specimens from patients enrolled in the EDALINE trial. The left panel is representative of a patient with high HPAS (characterized by high epithelial SHH ligand and high stromal GLI1 expression) while the right panel represents low/intermediate epithelial SHH and low stromal GLI1 expression. Scale bars: 100 µm. **C.** Representative immunohistochemistry staining for phospho-FGFR, collagen deposition depicted by SHG imaging and concomitant phospho-FAK and ALDH1 stem cell marker expression in patient tumor specimens with High Hh pathway activation signature from the EDALINE trial. The left panel represents the pre-treated biopsy from the patient who experienced a complete clinical response, the middle had a stable disease and the one on the right panel progressed on the prescribed regimen. Scale bars: 100 µm. **D.** Graphical summary: Paracrine Hh signaling in TNBC drives a reversible stem-like, drug-resistant phenotype *via* FGF signaling and ECM remodeling. CAFs represent the primary cells of the breast TME responding to Hh ligand. Hh-activated CAFs enhance ECM collagen deposition and express FGF5 in response to Hh signaling, which establish a supportive niche for cancer stem cell maintenance. This study strongly highlights a novel rational approach targeting both the tumor cells and their surrounding signaling support using SMO inhibitors in patients with metastatic TNBC.

To assess if stromal Hh pathway activation determines clinical response, we evaluated epithelial SHH ligand and stromal GLI1 expression in treatment naïve surgical tissue by immunochemistry. Only 10 patient tumor samples were evaluable. Three out of 10 tumors had high paracrine Hh Pathway Activation Signature (HPAS), characterized by high epithelial SHH in combination with high stromal GLI1 expression (**Fig. 7B** **and Supplementary Table S3**). Of these, two patients experienced a clinical response whereas all patients with low HPAS expression had progressive metastatic disease on the prescribed treatment regimen (**Supplementary Table S3**). An additional patient experienced clinical benefit, but the status of Hh pathway activation was unknown as her tumor sample was not available for analysis (**Supplementary Table S3)**.

Downstream analysis of the effect of paracrine Hh signaling revealed moderate to high phospho-FGFR expression, high collagen deposition and fiber linearization in treatment naive tumor specimens of the two responders with biopsy material available (**Fig. 7C**). This correlated with elevated phospho-FAK mechano-signaling and ALDH1 positive cells at the tumor-stromal interface (**Fig. 7C**). In contrast, the non-responder with high paracrine HPAS exhibited weak phospho-FGFR expression, low collagen content and minimal/no evidence of mechano-signaling and breast cancer stem cells (**Fig. 7C**). We therefore conclude that these additional tumor factors may represent adjunct biomarkers of therapeutic response for patient selection for anti-SMOi based combination therapies in Hh-expressing TNBC.

## DISCUSSION

In certain settings, CSCs are responsible for metastasis to distant organs (39-42) and are frequently enriched in residual tumors following chemotherapy (43-45), reflecting a role in therapeutic resistance. The hierarchical model for CSC maintenance proposes that CSC behave like tissue-resident physiological stem cells, self-renewing and undergoing asymmetric divisions to generate differentiated progeny (46). However, evidence from cell culture models has challenged the hierarchical CSC model, by suggesting that cancer cells can transition into a CSC state under specific culture conditions (46-49). In support of this notion, we now demonstrate that stromal cues from Hh-activated CAFs, forming a supportive niche enriched for FGF and fibrillar collagen-rich ECM, are capable of inducing and maintain a stem-like phenotype in TNBC cells *in vivo*. By combining a murine gain-of-function model, small molecule inhibitor studies in human xenografts with powerful *in vitro* systems, we have demonstrated the plastic characteristics of breast CSCs that can be successfully targeted using anti-stromal therapies, reducing metastatic growth and sensitizing to taxane chemotherapy.

Increased stromal collagen content correlates with stemness in the epidermis, both in the cancer and homeostatic contexts (50,51). It also enhances CSC properties of breast cancer cells *in vitro* (52,53). However, the impact of ECM collagen content and matrix mechanical properties on the biology of CSCs is not well defined. Our work provides new mechanistic insights, demonstrating that increased collagen density and fiber linearity at the tumor-stroma interface are associated with FAK activation and increased CSC number, dependent upon Hh paracrine signaling. Notably, we report a relationship between collagen abundance and clonogenicity *in vitro* and *in vivo*. Suppressing collagen production using SMO inhibitors was associated with decreased Krt6^+^ and ALDH1^+^ CSCs, respectively, in both murine and human models of TNBC. Interestingly, recent data links mammographic fibrillar collagen density to breast cancer risk, raising the possibility that breast cancer progenitors in these patients may have expanded in response to a dense collagen matrix (28,54,55).

FGF signaling has been shown to drive malignant processes including stem cell self-renewal, multipotency and therapeutic resistance (22,56,57). In metastatic breast cancer, resistance to anti-cancer treatment is primarily due to FGFR gene amplification (58). Here, we demonstrate a novel ligand-driven mechanism by which FGFR activation mediates both breast cancer stemness and chemoresistance, downstream of activation of the Hh signaling pathway. Importantly, our findings strongly suggest that CAF targeting using small molecule inhibitors of SMO is sufficient to prevent FGF ligand signaling and may overcome resistance to chemotherapies. Interestingly, FGF5 has been reported to be upregulated in prostate CAFs relative to normal fibroblasts (59), where it is also a target of Hh-Gli signaling (60). Thus this axis may be operational, and of therapeutic value, in tumor types beyond TNBC.

How FGFR activation and high FAK mechanosignaling lead to the establishment of a stem-like phenotype remains to be determined, but they are associated *in vitro* and *in vivo* with upregulation of transcription factors previously implicated in mammary physiological and cancer stem cells, including ID3 and SOX10 (21,22). The mechanisms underlying *Id3* and *Sox10* transcription are unknown. However *Id3* may be induced through Erk-EGR1 signaling, as observed in activated T cells, downstream of both FAK and FGFR (61). Our data also reveals the cooperative activity of ECM remodeling and FGF signaling in driving malignancy and drug resistance, recapitulating the interaction seen between these pathways during development and wound healing (62).

Importantly, many elements of Hh paracrine signaling to CAFs are active during embryonic development in mammals, though have not previously been linked. Dhh is highly expressed in a subset of epithelial cells of the mammary end bud, an invasive and proliferative structure responsible for ductal elongation in the developing mouse mammary gland (63). Consistent with our observation in TNBC, stromal but not epithelial Hh signaling is required for appropriate ductal morphogenesis (64). A number of FGF ligands are secreted by mammary stromal cells, and activation of epithelial FGFR1/2 is required for mammary ductal elongation and stem cell activity (65-67). In addition, mammary stromal fibroblasts secrete and remodel ECM components including collagens (68). Similar to our results in neoplastic cells, increased collagen density and mechanosignaling *via* FAK is sufficient to inhibit mammary epithelial cell differentiation and increased clonogenic potential (68). Thus the paracrine Hh signaling we observe in TNBC most likely represents the dysregulation and chronic activation of a process that is important for normal mammary ductal morphogenesis.

Using high-throughput single cell RNA-sequencing, we demonstrate that CAFs are the only stromal cell type responding to Hh ligand, and that SMO inhibitors act ‘on-target’ to reverse CAF gene expression changes induced by Hh signaling. Surprisingly, long-term (up to 3 months) daily treatment with SMOi did not alter the stromal cell composition of mammary tumors. This result contrasts markedly to that recently observed in pancreatic, colon and bladder cancer models, where chronic SMO inhibition was associated with marked changes in stromal cellular composition and shorter survival for mice receiving long-term SMOi treatment (69-72). The basis for this difference is not known, but may be explained by the divergent epi/genomics contexts of these cancer types, resulting in the evolution of distinct tumor microenvironments (11). Alternately, differences in the origin or phenotype of CAFs (73) in these endodermally-derived tumors versus ectodermally-derived mammary carcinomas may be relevant.

The benefit from therapeutic targeting of CAFs is two fold. Firstly, others and we have provided evidence for the crucial role of CAFs in supporting CSC self-renewal and resistance to chemotherapy (74-79). Therefore, targeting the CAF population and the subsequent abolition of the CAF-neoplastic cell interaction represent a practical strategy to improve cancer outcomes. Secondly, unlike neoplastic cells, CAFs have not been reported to exhibit genomic instability and are therefore less likely to acquire resistance to therapy over time, making them good targets for combination cancer therapies. Combined therapy with SMOi + docetaxel was well tolerated by mice and humans, and effective in treating a proportion of women with metastatic disease who had previously failed on taxane chemotherapy, including one patient who experienced a complete response. These remarkable results provide the first evidence to our knowledge for clinical benefit from a CAF-directed therapy. Treatment response in patients correlated with high levels of paracrine Hh signaling, FGFR activation and fibrillar collagen deposition, suggesting that the mechanism of action in patients may be consistent with that in mouse models. Hh, FGFR or collagen pathway activation may have value as predictive biomarkers of response to SMOi. Whilst phase I clinical trials are not designed nor powered to assess therapeutic efficacy, these data suggest an exciting new therapeutic strategy for drug-resistant or metastatic TNBC which should proceed to prospective assessment through Phase II clinical trials.

## METHODS

### Cell culture

M6 murine mammary carcinoma cells derived from the C3(1)/SV40 Tag mouse model (gift from J. Green, NIH (80)) were cultured as previously described (8). The human triple negative breast cancer cell line MDA-MB-231 was obtained from the American Type Cell Culture Collection (ATCC) and grown in RPMI-1640 supplemented with 10% (v/v) fetal bovine serum (FBS; Gibco^®^). All cell lines were grown at 37°C in a 5% CO_2_ incubator. Each cell line was characterized by short tandem repeat analysis (STR) profiling using the PowerPlexR 18D System (Promega) and tested for mycoplasma contamination (MycoAlert™ Mycoplasma Detection kit, Lonza). SMOi GDC-0449 (S1082, Selleckchem) and NVP-LDE225 (Novartis, Australia) were dissolved in DMSO to stock concentration of 10 mM. Docetaxel chemotherapy (McBeath, Australia) was diluted fresh (0.1 nM to 10 µM) with cell culture media. Recombinant FGF-5 (237-F5-050, R&D) was reconstituted at 10 µg/mL in PBS containing 1 µg/mL sodium heparin (H3149-50KU, Sigma-Aldrich) and 0.1% bovine serum albumin (A9418-10G, Sigma-Aldrich). NVP-BGJ 398, a potent and selective FGFR inhibitor (S2183, Selleckchem) was used at a concentration of 500 nM. All cell culture experiments involving SMOi (1 nM to 10 µM) and chemotherapy (IC30) lasted 5 days. For cell viability assays, treatment of M6 cells with recombinant FGF-5 (150 ng/mL) in the absence of FBS lasted 5 days. Cell viability assays were carried out in 96 well plates (Corning^®^ Life Sciences) and were determined by alamarBlue^®^ reduction.

### Tumor dissociation

M6 tumors were processed into single cell suspensions before limiting dilution assays, fluorescence-activated cell sorting (FACS) or flow cytometry analysis. Tumor dissociation into single cell suspension was carried out using the MACS mouse Tissue Dissociation Kit (Miltenyi Biotec, Australia) in gentleMACS C tubes on the gentleMACS Dissociator (Miltenyi Biotec) according to the manufacturer’s recommended protocol. Briefly, up to 1.5 g of tissue was transferred into a gentleMACS C-Tube (Miltenyi Biotec) containing 2.35 mL of RPMI 1640 solution with 1X Penicillin/Streptomycin (Gibco). 100 µL of enzyme D, 50 µL of enzyme R and 12.5 µL of enzyme A were then added and the sample was processed by running the defined gentleMACS program m_impTumor_02 on the gentleMACS Dissociator. The sample was incubated for 40 min at 37°C under continuous agitation and then processed using the gentleMACS program m_impTumour_03. The sample was resuspended in RPMI 1640 and filtered sequentially through 70 µM and 40 µM cell strainers (BD Falcon) and the resulting single cell suspension was centrifuged at 300 × g for 7 min. Cells were then resuspended in 1X BD Pharm Lyse™ lysing solution (555899, BD Biosciences) for 3 minutes at room temperature (RT) to lyse erythrocytes.

### Flow cytometry and FACS isolation

Cell sorting and flow cytometry experiments were performed at the Garvan Institute Flow Cytometry Facility. Flow cytometry was performed on a Becton Dickinson CantoII or LSRII SORP flow cytometer using BD FACSDIVA software, and the results were analyzed using Flowjo software (Tree Star Inc.). FACS experiments were performed on a FACS AriaII sorter using the BD FACSorter software.

Single-cell suspensions of primary M6 tumors were incubated with anti-CD16/CD32 antibody (1:200, BD Biosciences) in FACS buffer (PBS containing salts, 2% FBS, 2% Hepes) to block nonspecific antibody binding.

For the Next Generation Sequencing (NGS) experiment, single-cell suspensions were then pelleted and resuspended in FACS buffer containing the anti-EpCAM-PerCP/Cy5.5 (1:500, BioLegend^®^, Clone: G8.8) for 20 min on ice. Cells were then washed twice in FACS buffer before being resuspended in FACS buffer containing DAPI (1:1,000; Invitrogen) to discriminate dead cells. Stromal cells identified as DAPI^−^/GFP^−^/ EPCAM^−^ and M6 cancer cells identified as DAPI^−^/ GFP^+^/EPCAM^+^ were collected from at least 5 tumor specimens per treatment group.

For the isolation of M6-Ctrl or M6-Hh cells for the limiting dilution assays, cells were pelleted and resuspended in FACS buffer containing the following murine lineage markers (Lin^+^): anti-CD45-biotin (1:100; BD Pharmingen™, Clone: 30-F11); anti-CD31-biotin (1:40; BD Pharmingen™, Clone: 390), anti-TER119-biotin (1:80; BD Pharmingen™, Clone: TER119), and anti-BP1-biotin (1:50; Affymetrix eBiosciences, Clone: 6C3) for 20 min on ice. Cells were then pelleted and resuspended in FACS buffer containing streptavidin-APC-Cy™7 (1:400; BD Pharmingen™) and the anti-EpCAM-PerCp/Cy5.5 (1:500, BioLegend^®^, Clone: G8.8), and incubated for 20 min on ice. Cells were then washed twice in FACS buffer before being resuspended in FACS buffer containing DAPI (1:1,000; Invitrogen). Live M6 primary neoplastic cells were sorted based on their GFP^+^/EpCAM^+^/Lin^−^ expression.

For the analysis of CSC properties, single-cell suspensions from M6 tumor models were incubated with the combination of the following murine lineage markers: anti-CD31-biotin (1:40), anti-CD45-biotin (1:100), anti-TER119-biotin (1:80), and anti-BP1-biotin (1:50) for 20 min on ice. Cells were then pelleted and resuspended in FACS buffer containing streptavidin-APC-Cy™7 (1:400) and the following epithelial stem cell markers: anti-CD24-PE (1:500; BD Pharmingen™, Clone: M1/69), anti-CD29-APC-Cy7 (1:100; BioLegend, Clone: HMβ1-1) and anti-CD61-APC (1:50; ThermoFisher), and incubated for 20 min on ice. Cells were then washed twice in FACS buffer before being resuspended in FACS buffer containing DAPI (1:1,000; Invitrogen).

### Co-culture of primary cells

Primary M6 cells and cancer associated fibroblasts (CAFs) derived from M6-Ctrl and M6-Hh tumors, respectively, were selected by FACS using the following cell surface markers: Epithelial cells: CD45^−^/CD31^−^/CD140a^−^/GP38^−^/EpCAM^+^/GFP^+^; CAFs: CD45^−^/CD31^−^/CD140a^+^/GP38^+^/EpCAM^−^/GFP^−^. After tumor dissociation, single cells were pelleted and resuspended in FACS buffer containing the following antibodies: anti-CD45-APC-eFluor^®^-780 (1:500; Affymetrix eBioscience, Clone: 30-F11), anti-CD31-biotin (1:100; BD Pharmingen™, Clone: 390), anti-CD140a-APC (1:100; BioLegend^®^, Clone: APA5), anti-Podoplanin-PE (1: 1,000, BioLegend^®^, Clone: 8.1.1) for 20 min on ice. After two washes with PBS, cells were then pelleted and resuspended in FACS buffer containing streptavidin-APC-Cy™7 (1:400; BD Biosciences), and incubated for 20 min on ice. Cells were then washed twice in FACS buffer before being resuspended in FACS buffer containing DAPI (1:1,000; Invitrogen) to discriminate dead cells. Different epithelial and CAF populations from at least 3 tumor specimens per treatment group were isolated and cultured into 100 mm culture dishes (Corning^®^ LifeSciences) in DMEM supplemented with 10% (v/v) FBS, 50 µg/mL gentamycin and 1x antibiotic/antimycotic (15-240-096, Gibco^®^) in a 5% O_2_, 5% CO_2_ incubator at 37°C. 10^5^ M6 cells were added to the petri dish when CAFs derived from the corresponding tumors have reached 70% confluency and the co-culture assays lasted a total of 5 days.

### Animals and surgery

All animal procedures were carried out in accordance with relevant national and international guidelines and according to the animal protocol approved by the Garvan/St Vincent’s Animal Ethics Committee (Animal ethics number 11/46). M6-Ctrl or M6-Hh (0.75 × 10^6^ cells/10 µL) and MDA-MB-231 (1 × 10^6^ cells/10 µL) transplants were carried out by surgical injection via direct visualization into the fourth mammary fat pads of pre-pubescent Rag^−^/^−^ and NOD-scid IL2rγnull (NSG) mice, respectively.

For limiting dilution studies, single-cell suspensions of viable M6-Ctrl or M6-Hh tumor cells were prepared as described in the “Tumor dissociation” section. EpCAM^+^/GFP^+^/Lin^−^ tumor cells, isolated by FACS, were transplanted in appropriate numbers into the fourth mammary fat pad of 3 to 4-week-old syngeneic Rag^-/-^ mice and aged till ethical end point. Extreme limiting dilution analysis (ELDA (23)) software was used to calculate the tumor-propagating cell (TPC) frequency.

PDX tumor tissues, acquired from the laboratory of A. Welm (35) were serially passaged as 2mm^3^ fragments in the cleared fourth mammary fat pads of pre-pubescent NSG mice according to established protocols (35). When tumors became palpable, they were measured three times weekly in a blinded manner using electronic calipers to monitor growth kinetics. Tumor volume was calculated using the formula (π/6) × length × width^2^. Upon reaching ethical or predefined experimental endpoints, mice were euthanized and primary tumor and any associated metastases were collected.

### *In vivo* drug treatment experiments

SMOi GDC-0449 (S1082, Selleckchem) and NVP-LDE225 (Novartis, Australia) were dissolved in 0.5% methylcellulose, 0.2% Tween^®^ 80 (Sigma-Aldrich) and 0.5% methylcellulose, 0.5% Tween^®^ 80, respectively, and then delivered by oral gavage (100 mg/kg/bid, GDC-0449; 80 mg/kg/day, NVP-LDE225). Chemotherapy (Docetaxel, McBeath Australia) was diluted in 5% dextrose then delivered by intraperitoneal injection (15 mg/kg/week). Tumor bearing mice were randomly assigned into respective treatment groups once tumor volume reached 100 mm^3^ (*n* = 7 – 8 mice per group). Tumor growth was calculated for each individual tumor by normalizing to the tumor volume at day 0. In short-term studies examining the molecular and histological impact of Hh pathway activation and inhibition, mice were treated between 8 to 14 days then euthanized. At euthanasia, primary tumors were harvested and macroscopic metastatic lesions were scored. For the long-term therapeutic study, mice were treated to endpoint. Animals were excluded from overall survival analysis if they had to be sacrificed for poor body conditioning, unrelated to tumor size endpoint. Animal technicians, who were blinded to the experiment treatment groups, independently monitored the mice.

### Next Generation Sequencing

We isolated by FACS the stromal DAPI^−^/GFP^−^/EPCAM^−^ and epithelial DAPI^−^/ GFP^+^/EPCAM^+^ cell fractions from at least 5 M6-Ctrl and M6-Hh tumor models treated with vehicle (0.5% methylcellulose, 0.2% Tween^®^ 80) or with SMOi. RNA was isolated using the miRNeasy kit (Qiagen). For standard input samples, 1 µg of total RNA was used as input to the TruSeq RNA Sample Preparation Kit v2 (Illumina). The samples were prepared according to the manufacturer’s instructions, starting with the poly-A pulldown. The number of PCR cycles was reduced from 15 to 13, to minimize duplications. The samples were sequenced on the HiSeq2000 using v3 SBS reagents (Ramaciotti Centre for Genomics, University Of New South Wales (UNSW)). Low input RNA stromal samples were firstly amplified using the Ovation^®^ RNA-Seq System V2 kit (4 ng of total RNA input; Nugen Integrated Sciences Pty. Ltd.) according to the manufacturer instructions. 1 µg of the cDNA was sheared with Covaris to fragment sizes of ~200bp. The material was used as input to the TruSeq RNA Sample Preparation v2 kit, starting at the end-repair step. The number of PCR cycles was reduced from 15 to 10. All the samples were sequenced on the HiSeq2000 using v3 SBS reagents (Ramaciotti Centre for Genomics, University Of New South Wales).

### Bioinformatics and computational analysis of RNA-sequencing data sets

Analysis of the RNA-Sequencing data was conducted on the high-performance computing cluster at the Garvan Institute following a standard four step approach, cleaning, aligning, counting and differential expression with an additional normalization step. FASTQ files were quality checked using FastQC version 0.11.1 (<u>http://www.bioinformatics.babraham.ac.uk/projects/fastqc/</u>) and FastQ Screen version 0.4.4 (<u>http://www.bioinformatics.babraham.ac.uk/projects/fastq_screen/</u>) then quality filtered using FastqMCF version 1.1.2 (<u>https://code.google.com/p/eautils/wiki/FastqMcf</u>) to remove low quality bases and adapter contamination. Filtered reads were then aligned to the mouse reference genome GRCm38/mm10 using STAR, version 2.4.0d (Dobin, 2013). Feature counting was performed using HTSeq version 0.5.4p3 (Anders, 2014). Due to high levels of variation in the expression data between replicates, the RUV normalization procedure was implemented (81). This aims to remove unwanted variation and produce more reliable pair-wise comparisons when calculating differential expression. In this instance, RUVr with a K of 3 was found to be the most effective method based on the suggested diagnostics, e.g. plots of P-value distributions and PCA. Differential expression analysis was performed within the RUV analysis using edgeR (82).

### Single cell RNA-Sequencing using the Chromium Platform and bioinformatical analysis

M6 tumors were processed into single cell suspensions as described previously. Sorted live single epithelial and stromal cells were loaded on a Chromium Single Cell Instrument (10x Genomics, Pleasanton, CA) to generate single cell GEMs. As per manufacturer instructions, approximately ~ 7,000 cells were loaded per channel for a target of ~ 4,000 cells (10x Genomics). Two biological replicates were analysed per sample. Single cell RNA-Seq libraries were prepared using the Chromium Single Cell 30 Gel Bead and Library Kit (10x Genomics). Single-cell sequencing libraries were generated from sorted cells collected in parallel on the same Chromium Single Cell Chip and sequenced in multiplexed pairs to minimize experimental variability and any confounding batch effects. Single cell libraries were sequenced using the Illumina NextSeq 500 system using the following parameters: pair-end sequencing with dual indexing, 26 cycles for Read1, 8 cycles for I7 Index Read and 98 cycles for Read2.

The Cell Ranger Single Cell Software v2.0 (10X Genomics) was used to process raw bcl files to perform sample demultiplexing, barcode processing and single cell 3’ gene counting (<u>https://software.10xgenomics.com/single-cell/overview/welcome</u>). Reads were mapped to the mm10 mouse reference genome. The Cell Ranger aggregation pipeline was used to normalize the sequencing depths of multiple datasets (based on the proportion of mapped reads) to re-compute a combined gene-cell barcode matrix.

Downstream filtering and analysis of the raw gene-cell barcode matrix was performed using the Seurat v2.0 package in R (83). The gene-cell barcode matrix was filtered based on number of genes detected per cell (any cells with less than 500 or more than 6000 genes per cell were filtered), a total number of unique molecular identifiers (UMIs) (any cells with UMIs > 50,000 were filtered) and a percentage of mitochondrial UMI counts (any cells with more than 10% of mitochondrial UMI counts were filtered). Altogether, in the combined 5 datasets, a total of 14,950 cells and 18,487 genes were analyzed. Based on an expression cut-off of 0.0125 and a dispersion cut-off of 0.5, a total of 2,620 variable genes were selected for principal component analysis (PCA). A total of 91 significant principal components (determined using JackStraw in Seurat, *P* < 0.05) were used for clustering analysis and *t*-SNE projection. The classification of cell clusters was inferred using the following canonical markers: CAFs (*Pdpn* and *Pdgfrb)*, epithelial cancer cells (*Epcam),* endothelial cells (*Cd34* and *Pecam1*), macrophages/monocytes (*Ptprc* and *Cd68*), neutrophils (*Ptprc* and *Csf3r*) and natural killer cells (*Ptprc* and *Ncr1)*. Differential gene expression analysis in Seurat was performed using the ‘bimod’ likelihood-ratio test.

### Data and Code Availability

All RNA Sequencing files that support the findings of this study have been deposited in GEO with the accession code PRJNA369574. The RNA-Seq pipeline and the analysis scripts can be found on the respective websites: <u>https://github.com/elswob/rna-seq-pipe</u> and <u>https://github.com/elswob/Hh</u>.

### Gene Set Enrichment Analysis (GSEA)

Gene-sets used in GSEA were extracted from version 3.1 and 4.0 of the Broad institute’s Molecular Signatures Database (MSigDB) (84,85) and extended with additional curated gene-sets from literature. All GSEA analyses were performed using a combined set of the c2, c5 and c6 from MSigDB plus additional curated sets that we identified in the literature.

### Association of the Hedgehog stromal gene signature with clinical outcome using TCGA expression data

A stringent mouse gene signature was derived through differential expression analysis of the stromal fraction of M6-Hh tumors in comparison to the stroma of M6-Ctrl or M6-Hh tumors treated with SMOi. The stromal mouse gene signature was then converted to human genes using NCBI homolog gene list v68 (<u>http://www.ncbi.nlm.nih.gov/homologene</u>). The converted gene list consists of 146 upregulated genes and 39 downregulated genes in the Hh-activated stromal population (**Supplementary Table S1**). The gene list was further assessed for survival analysis using the TCGA breast invasive carcinoma cohort. The processed TCGA data was downloaded from cBioPortal (86) based on the TCGA study (87). The gene signature score was defined by a weighted average method for each sample in the TCGA cohort. Survival curves were estimated using the Kaplan-Meier method, with overall survival used as the outcome metric.

### RNA isolation, reverse transcription, quantitative RT-PCR and Fluidigm array experiments and analysis

Individual stromal CAFs and epithelial malignant M6 cancer cells were FACS-isolated as described above. For each fraction, between 1,000 and 50,000 cells were directly collected into QIAzol lysis reagent. Low input RNA samples were then isolated using the miRNeasy Micro kit (Qiagen). All the other standard input RNA samples were isolated using the miRNeasy kit (Qiagen) and reverse transcribed with the Transcriptor First Strand cDNA Synthesis Kit (Roche Diagnostics). cDNA was synthesized from 0.5 – 2 µg of total RNA and diluted 1:10 before any further quantitative RT-PCR (qRT-PCR) analysis.

qRT-PCR experiments were performed using either the Roche Universal Probe Library System on a Roche LightCycler480^®^ (Roche LifeScience) or the TaqMan Gene Expression Assay (Applied Biosystems/ Life Technologies) on an ABI Prism^®^ 7900 HT Sequence Detection System (Perkin-Elmer Applied Biosystems). Primers, probes and programs used for qRT-PCR analysis are listed in **Supplementary Table S5**. Relative mRNA expression levels were normalized to β-actin, GAPDH or HPRT and quantification was performed using the comparative C_T_ method described by Livak and Schmittgen (88).

### Immunohistochemistry, immunofluorescence and histological analysis

Tissues were fixed in 10% neutral buffered formalin at 4°C overnight then processed for paraffin embedding. For histological analysis, 4 µm tissue sections were stained with haematoxylin and eosin using standard methods. Immunohistochemical, immunofluorescence and picrosirius red staining were performed on paraffin-embedded tissue sections using standard protocol. Full details of each antibody used and their relative staining protocols for immunochemistry are described in **Supplementary table S6**.

Histological analysis of the proliferative marker phospho-histone H3, ALDH-1 and the progenitor cell marker CK6 were carried out by digitizing entire images using the Aperio CS2 digital pathology slide scanner (Leica Biosystem) at 20x magnification. Cells that stained positively for phospho-Histone H3, ALDH-1 or CK6 within a distance of 200 µm from the CAFs at the tumor-stromal interface were then counted and averaged over at least 5 fields using the Aperio Imagescope software (Leica Biosystem). The limit of 200 µm reflects the well-established diffusional distance for Hh ligand in mammalian models (89). Picrosirius red stain was analyzed as previously described (90). Two specialist breast pathologists, who were blinded to the experiment treatment groups, independently scored the remaining IHC stains. Areas of necrosis were excluded for all analysis. The number of CD45^+^ cells in the peritumoral stroma and the number of CD31^+^ blood vessels w estimated and averaged over 5 high power fields (40x magnification). Alpha-SMA was scored as the percentage of myofibroblabts in the tumor stroma at the whole tumor level. Tumor versus stromal ratio was estimated on H&E sections.

For the immunofluorescence staining, formalin-fixed paraffin-embedded sections on Superfrost+ glass slides (Thermo) were dewaxed and rehydrated, and antigen retrieval was performed by boiling for 12 minutes in 10 mM Citrate buffer (pH 6.0) or 10 mM Tris-Cl (pH 9.0) in a pressure cooker (**Supplementary Table S6**). Following blocking with 10% goat serum, sections were incubated with the primary antibodies (p(Tyr397)-FAK, CK6, ALDH1, E-cadherin at 4°C for 18 hours, washed in PBS and incubated with the appropriate secondary antibodies for 1 h. After washing in PBS, sections were incubated with 0.2 µg/ml DAPI in PBS for 5 min to label nuclei and mounted. Confocal fluorescence images were captured using a LSM 700 confocal scanning system (Carl Zeiss AV), with Zen 2011 (Black Edition) version 8.1.5.484 software. Images were processed using ImageJ (National Institutes of Health) as previously described (51,91). Two cell biologists from various institutes, who were blinded to the treatment groups, independently scored the IF stains for CK6, ALDH1 and phospho-FAK.

### Second-harmonic generation (SHG) microscopy, Gray-level co-occurrence matrix and orientation analysis

Formalin-fixed, paraffin embedded sections stained with hematoxylin and eosin and mounted in DPX (Sigma) were imaged using a 20x 1.0 NA objective on an upright fixed-stage two-photon laser scanning microscope system (Zeiss). The excitation source was a Ti:Sapphire femto-second laser cavity (Newport Mai Tai), coupled into a LSM 710 scan module. An excitation wavelength of 890 nm was used to collect SHG signal (435 ± 20 nm) from collagen. Maximum collagen coverage values derived from SHG signal (by depth (line graph) and at peak value (histogram inset)) was used as a measure of collagen abundance and density. Signal was acquired from three separate areas measuring 320 × 320 µm^2^ across each sample. Bright-field transmission images were co-acquired with SHG data.

ImageJ (NIH, Bethesda MD, USA) was used to calculate percentage area covered by SHG signal per image, after conversion to a binary image based upon a single manually determined threshold value applied across all images as previously described (50,91). Results were expressed as medians, ranges and quartiles across all data sets.

Gray-level co-occurrence matrix (GLCM) analysis was carried out as previously described (31). Briefly, collagen fiber organization was assessed using GLCM analysis to characterize the texture of a sample and determine the correlation of the SHG signal within the matrix. The correlation plots represent the similarity in signal strength between pixels. A slower decay shows a more organized and correlated network of collagen fibers than in samples with a faster decay. GLCM analysis was performed in ImageJ.

Orientation Analysis was carried out as previously described (30). Briefly, fiber orientation analysis was performed on SHG images using an in-house ImageJ (NIH) macro where structure tensors were derived from the local orientation and isotropic properties of pixels that make up collagen fibrils. Within each input image, these tensors were evaluated for each pixel by computing the continuous spatial derivatives in the x and y dimensions using a cubic B-spline interpolation. From this, the local predominant orientation was obtained. The peak alignment (measured in degrees) of fibers was then determined, and the frequency of fiber alignment calculated.

### Alginate-Collagen I Inter-Penetrating Network (IPN) hydrogels

Alginate IPNs were generated from 1% sodium alginate mixed with either 20%, 5% or 0% rat tail collagen I, plus 1% Matrigel. M6-Ctrl and M6-Hh single cells were encapsulated in 400 µm diameter beads before transferring to normal growth media. Colony forming ability was assessed at day 5, 8 and 12.

### Tumorsphere assays

Low passage M6 cells grown to 70 – 80% confluency as adherent monolayer were trypsinized, quenched in normal culture media, washed three times with large volumes of calcium-magnesium free PBS then passed through a 40 µm cell strainer to obtain a single cell suspension. Cell number was determined using the Countess™ Automated Cell Counter (Invitrogen) then seeded in sphere-promoting culture at density of 2.5 × 10^3^ cells/mL in ultra low-adherent 6-well plates (Corning^®^ LifeSciences). Cells were grown at 37°C in a 5% CO_2_ incubator. Primary sphere formation efficiency was determined after 5 days. Spheres larger than 40 µm were counted manually using a light microscope and automatically using the IncuCyte ZOOM^®^ Live Cell System (Essen BioScience). Primary spheres were then collected by gentle centrifugation and washed with calcium-magnesium free PBS prior to dissociation into single cell suspension. Cell number was determined as above then seeded in triplicate at density of 5 × 10^2^ cells/mL in ultra low-adherent 6-well plates. Secondary sphere formation efficiency was determined after 8 days. Sphere media was composed of DMEM/F12, 1% methylcellulose, 1 x B27 supplement (17504-044, Invitrogen), and 4 µg/mL sodium heparin (H3149-50KU, Sigma-Aldrich). Tumorspheres were treated with 150 ng/mL recombinant FGF5 (237-F5-050, R&D Systems) and / or 500 nM FGFR inhibitor, NVP-BGJ 398 (S2183, Selleckchem). Sphere formation efficiency was calculated using the following formula: (Number of tumorspheres larger than 40 µm / Number of single cells seeded) x 100%.

### Patients

Patients in this study were enrolled in the GEICAM/2012-12 (EDALINE) clinical trial (ClinicalTrials.gov Identifier: NCT02027376); a single-arm, open-label, phase I, 3+3 dose escalation study in which patients with TNBC were treated with the SMOi Sonidegib (LDE-225) in combination with docetaxel to determine the Maximum Tolerated Dose and the Recommended Phase II Dose (RP2D), as the primary objective. Written informed consent was obtained from all patients and documented before performing any protocol-specific procedure. The study was conducted in accordance with the International Conference on Harmonization Good Clinical Practice Guidelines (ICH GCP), the Declaration of Helsinki and applicable local regulatory requirements and laws. The protocol was approved by the Institutional Review Board and the Ethics Committee of all the participating sites (Hospital General Universitario Gregorio Marañón, Hospital Universitario Clínico San Carlos, Complejo Hospitalario Universitario A Coruña, Hospital Universitario Virgen del Rocío, Hospital Universitario Virgen de la Victoria), according to the requirements of the Spanish regulations (GEICAM/2012-12; clinicaltrials.gov identifier: NCT02027376). Eligible patients, with no more than three previous lines of chemotherapy for metastatic disease were treated with 21-day cycles of intravenous docetaxel 75 mg/m^2^ on day 1 and oral Sonidegib administered at increasing doses of 400, 600 and 800 mg once daily (QD), until radiographic or symptomatic progression, unacceptable toxicity or withdrawal of the informed consent, whichever occurred first. All patients at each dose level completed at least 2 treatment cycles before being enrolled to the next Sonidegib dose level. Twelve patients were enrolled into the study and were treated as described above.

### Clinical markers of therapeutic activity

The expression of Sonic Hedgehog ligand (SHH) and GLI1 were examined by immunohistochemistry (IHC) from archived formalin-fixed paraffin-embedded (FFPE) pre-treatment primary breast tumors from patients enrolled in the EDALINE clinical trial. Pathologists, who were blinded to clinical parameters, carried out biomarker analysis. Two specialist breast pathologists independently calculated the Histo-score (Hscore) based on the percentage of stained cells (SHH and GLI1 expression) and staining intensity on a predetermined scale (0: no staining to 3: strong), in the tumor epithelial cells (SHH) and the tumor stroma (GLI1). Predefined cut-offs for high/low biomarker expression were established based on standard criteria (median Hscore for SHH in tumor cells and intensity staining for GLI1 in the stroma).

We defined a Hedgehog Pathway Activation Signature (HPAS) predictive for clinical response to sonidegib (LDE-225) in combination with docetaxel as cases with both high SHH expression in the tumor epithelium (SHH Hscore > 150) and intense GLI1 expression in the tumor stroma.

### Statistical analysis

All statistical analyses of all data were performed using GraphPad Prism 6.0c software (GraphPad Software). For all *in vitro* experiments, three or six technical replicates were analyzed for each experiment, and results are presented as the mean ± s.e.m. of three biological replicates. Quantitative analyses were carried out using unpaired two-tailed Student’s *t*-test with equal standard deviation after confirming that the data met appropriate assumptions (normality, homogenous variance, and independent sampling). 7-10 mice per treatment group were used for all *in vivo* experiments with SMOi treatment. 5 mice per treatment group were utilized for the RNA-Seq. For all RT-qPCR experiments, three technical replicates were analyzed for each experiment, and results are presented as the mean ± s.e.m. of three biological replicates. Subsequent statistical analysis from *in vivo* experiments was performed with either unpaired two-sided student *t*-tests or the Fisher’s exact test. Survival analysis was performed using the Log-rank (Mantel-Cox) test. A *P* value < 0.05 was considered statistically significant. \**P* < 0.05, \*\**P* < 0.01, \*\*\**P* < 0.001, \*\*\*\**P* < 0.0001. Estimation of variation within each group was determined by s.e.m. Sample size estimation was initially chosen by using power calculations for guidance (<u>http://biomath.info/power/ttest.htm</u>). Effect sizes were estimated at 30% for single agent arms, based on earlier data with single agent docetaxel and SMO inhibitors.

With alpha = 0.05 and power = 0.9 and allowing for attrition of ~ 1 mouse/group, ≥ 7 mice per group were needed for all preclinical studies. For the Phase I clinical trial EDALINE, descriptive analysis on demographic and clinicopathological characteristics (age, visceral disease, number of involved sites, prior treatment, histologic tumor grade and Ki67), Hh biomarkers expression (SHH and GLI1) and efficacy data to the combined therapy sonidegib with docetaxel were performed. The Chi-Square Test of Independence was assessed to examine the association between epithelial SHH and stromal GLI1 expression and efficacy endpoints: best tumor response (complete or partial response, stable disease, progression disease and Time to Progression (TTP)). TTP was defined as the time from treatment commencement until objective tumor progression (does not include deaths). Progression-Free Survival (PFS) was explored using Kaplan-Meier survival analysis. PFS was defined as the time from treatment commencement until disease progression or death. For mouse and clinical studies, specialists were blinded to the experiment treatment groups.

## AUTHORS CONTRIBUTION

### Conception and design

Aurélie S. Cazet, Mun N. Hui, Sandra O’Toole, Miguel Martín, Alexander Swarbrick

### Development of methodology

Aurélie S. Cazet, Mun N. Hui, Benjamin L. Elsworth, Sunny Z. Wu, Daniel Roden, Thomas R. Cox, Michael Samuel

### Acquisition of data (provided animals, acquired and managed patients, provided facilities, etc.)

Aurélie S. Cazet, Mun N. Hui, Sunny Z. Wu, Chia-Ling Chan, Joanna Skhinas, Raphaël Collot, Jessica Yang, Kate Harvey, M. Zahied Johan, Caroline Cooper, Radhika Nair, David Herrmann, Michael Samuel

### Analysis and interpretation of data (e.g., statistical analysis, biostatistics, computational analysis)

Aurélie S. Cazet, Mun N. Hui, Benjamin L. Elsworth, Sunny Z. Wu, Daniel Roden, Nian-Tao Deng, Thomas R. Cox, Michael Samuel, Alexander Swarbrick

### Writing, review, and/or revision of the manuscript

Aurélie S. Cazet, Mun N. Hui, Elgene Lim, Paul Timpson, Sandra O’Toole, D. Neil Watkins, Thomas R. Cox, Michael Samuel, Miguel Martín, Alexander Swarbrick

### Administrative, technical, or material support

Jessica Yang, Kate Harvey, Andrea McFarland

### Other (clinical trial management, oversight of trial conduct, and sample collection)

Manuel Ruiz-Borrego, Federico Rojo, José M. Trigo, Susana Bezares, Rosalía Caballero

### Study supervision

Miguel Martín, Alexander Swarbrick

## ACKNOWLEDGMENTS

We would like to thank the following people for their assistance with this manuscript; Ms. Nicola Foreman and Ms. Breanna Fitzpatrick for animal handling, Ms. Alice Boulghourjian and Ms. Anaiis Zaratzian for tissue processing and IHC staining, Mr. Rob Salomon, Mr. Eric Lam, Mr. David Snowden and Mrs. Vitri Dewi for their help with flow sorting, Ms. Natasha T. Pyne for SHG imaging, Ms. Stephanie Ketterer, Ms. Sylvia Pietkiewicz and Mrs. Natahsa Sharma. Mr Julien Charton for his help with the proofreading.

We are grateful to the patients and families involved in this study.

